# Experimental design and analysis of sulfide-induced invasive growth in wine yeast

**DOI:** 10.64898/2026.04.06.716814

**Authors:** Kai Li, Jennifer M. Gardner, Lauren Kennedy, Jin Zhang, Joanna F. Sundstrom, Stephen G. Oliver, Alexander K. Y. Tam, J. Edward F. Green, Vladimir Jiranek, Benjamin J. Binder

## Abstract

Yeast’s ability to invade surfaces has important implications for infections and food contamination. Invasive growth in yeast is influenced by genetic and environmental factors. In this exploratory study, we used a systematic experimental design to identify conditions under which sulfide-induced invasive growth can be reliably observed and quantified and investigated the effects of sulfide, gene deletions, and environmental conditions on the invasive behaviour of the wine yeast strain AWRI 796. Sulfide enhanced invasion in the (parent) AWRI 796 strain under nitrogen-limiting conditions, although its effect was obscured by experimental variability and pre-culture conditions. Genetic factors had a major effect on the overall invasive phenotype, with deletion of key genes suppressing invasion. Most gene-deletion mutants did not significantly affect how the colony responded to sulfide. In addition to sulfide and genotype, environmental conditions also influenced invasive behaviour. The pre-2×SLAD pre-culture condition was best for detecting sulfide-induced growth, and later plate washing time and decreased nutrient levels enhanced invasiveness. Our experimental design and findings provide a framework for understanding the determinants of yeast invasiveness, which may inform future studies on filamentous yeast behaviour.

## Introduction

The budding yeast *Saccharomyces cerevisiae* is well known for its role in baking and brewing, and is also an important model organism in scientific research. This yeast can change its mode of growth in response to challenging environments^1,2^. When nutrients become scarce or conditions become stressful, for example due to the lack of nitrogen, the presence of some aromatic alcohols, or changes in acidity, yeast cells can shift from growing as individual rounded cells to forming chains of elongated cells that spread across and into surfaces^3–5^. These chains are are not true hyphae, and are instead known as pseudohyphal filaments, which consist of connected individual cells that remain attached after division^3,6–8^. Pseudohyphal filamentous growth allows yeast communities to forage and attach to substrates more effectively, enabling access to more favourable environmental conditions such as new food sources^3,9–11^. Since both grape juice and oak sap are known to be nitrogen poor^12,13^, pseudohyphal growth may be relevant to the yeast’s lifestyle in industry and in the wild. Pseudohyphal growth may also enable *S. cerevisiae* to adhere to or penetrate plant tissues, providing access to new sources of nitrogen^8^.

In laboratory experiments, pseudohyphal filamentous growth in *S. cerevisiae* on solid substrates (most commonly agar based media) can be observed in two closely related but distinct ways^3,4^. Under certain conditions, the colony produces visible pseudohyphal filaments at the colony edge. Pseudohyphal filaments can also grow downward into the agar, which is termed invasive growth. Both surface spreading and invasion are commonly triggered by the aforementioned environmental stresses such as nitrogen (*e*.*g*. ammonium) limitation, and can be readily studied using simple plate-based assays (Figure 1). The combination of filamentous surface growth and invasive growth indicates how yeast respond to their environment^8,10^.

**Figure 1.**
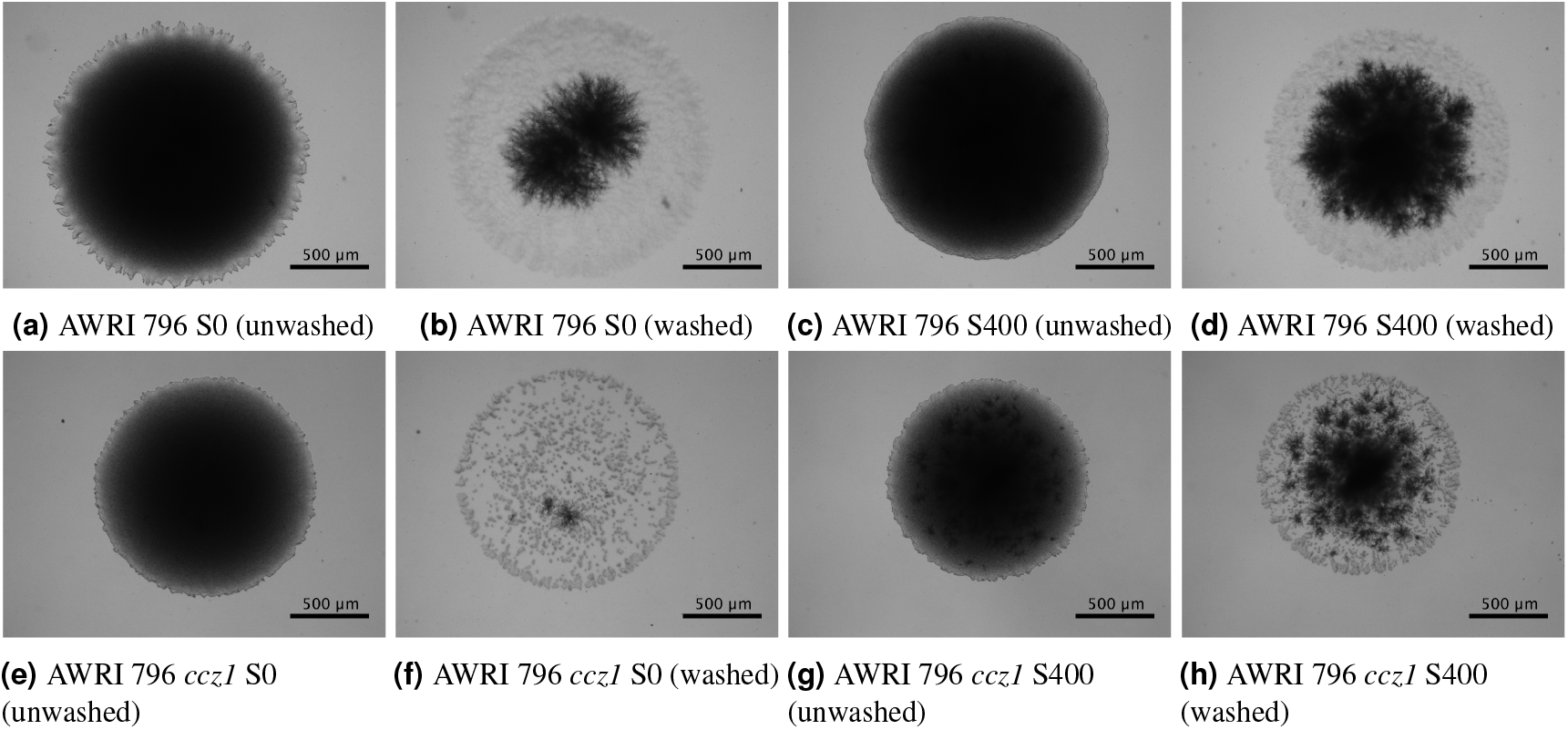
Colony images of AWRI 796 and a gene-deletion mutant (AWRI 796 *ccz1*) of unwashed (capturing the surface growth a, c, e, g) and washed (revealing invasively growing cells b, d, f, h) colonies at day 6 without (S0) and with (S400) sodium sulfide (400 µM).

Filamentous surface growth is readily observed in top-down images of lab-grown colonies (first and third columns Figure 1). In contrast, invasive growth is an inherently three-dimensional process that occurs beneath the agar surface. Unlike surface growth, invasive growth cannot be observed using standard widefield light microscopy, and requires complex microscopy to capture in detail^14^. Due to this limitation, invasive growth is often assessed using a simpler indirect assay. This involves washing the plate to remove non-adherent surface cells, leaving behind only those cells that have penetrated into or are strongly adhered to the agar (second and fourth columns Figure 1). The two-dimensional pattern that remains on the plate surface then indicates the extent of invasive growth^15^. Quantifying invasive growth in these indirect assays presents practical challenges. Growth into the medium can only be assessed after the surface colony has been removed. However, if plates are washed too late, it can be difficult to detect meaningful differences because many cells within a colony may grow invasively regardless of condition. Conversely, washing too early may underestimate invasion because insufficient time has elapsed for invasive growth to occur.

Given the difficulties of quantifying invasive growth, we considered two complementary measures, the *presence of invasion* and the *degree of invasion*. The presence of invasion is defined as the proportion of colonies on a plate that show any visible invasive growth after washing. This measure is calculated as the number of colonies displaying invasion divided by the total number of colonies on the plate. Since all four colonies in Figure 1 invade, in this example the presence of invasion would be one. The presence of invasion can be assessed reliably and quickly by visual inspection, using all colonies on the plate. The degree of invasion measure is computed for individual colonies. For a given colony, the degree of invasion is defined as the washed (invasive) area divided by the unwashed (surface) area. Computing the degree of invasion requires processing close-up images of a colony, which may only be available for a subset of the colonies. We used our custom-built, open-source software, TAMMiCol, to convert colony images into binary images as an initial step^16^. Additional processing was then applied to distinguish the darker invasive growth from the lighter, non-invasive areas of the colony (see Figure 1). We then used the resulting processed binary images to calculate the degree of invasion for each available colony.

Of the environmental stresses known to promote invasive growth, we focused on how sulfide exposure affects invasive growth in *S. cerevisiae*. Yeasts produce sulfides in natural and industrial environments, particularly when starved of nitrogen^17–19^. However, the effect of sulfides on surface filamentous and invasive growth remains underexplored. Additionally, sulfide has also been proposed to function as a signalling molecule in cellular processes, where it can influence gene regulation and stress responses^20–25^. This raises the possibility that sulfide may actively promote invasive growth through specific biological pathways. To study this possibility further, we also examined the impact of exogenously added sulfide on invasive growth of a set of key gene-deletion mutants. These were homozygous deletions constructed in the diploid AWRI 796. We targeted genes that have already been reported to be involved in yeast sulfur metabolism. Genes were selected from those known to function within an associated biochemical pathway or phenotype, for instance “Sulfate assimilation pathway” or “utilisation of sulfur source absent” according to the annotation tools in the *Saccharomyces* Genome Database^26^, and relevant literature mining (Supplementary material, Table 3). Our manually curated list was then filtered for genes that encoded proteins likely to be transporters, in an effort to discover candidate genes that may function as sulfide sensors, and thus potentially trigger invasive growth. We also filtered for genes known to be involved in surface filamentous and/or invasive growth and other associated phenotypes such as genes that result in cell elongation or hyperpolarised growth when overexpressed. Many of these genes play roles in environmental sensing, stress responses, and growth regulation, making them plausible candidates for mediating the effects of sulfide^27^.

The primary aim of this work was to use a systematic experimental design to identify conditions under which sulfide-induced invasive growth can be reliably observed, while also investigating the effect of sulfide and genetic background on invasive growth. Figure 1 compares the invasive growth in the absence and presence of sulfide. Notably, greater invasive growth occurred when sulfide was present in both AWRI 796 and its gene-deletion mutant AWRI 796 *ccz1*. In the absence of sulfide, the gene-deletion mutant AWRI 796 *ccz1* exhibited noticeably less invasive growth than the parent AWRI 796 strain (second column). When sulfide was added, both the parent and the mutant showed increased invasion, with the increase appearing more pronounced in the mutant (fourth column). This apparent interaction suggests that deletion of *ccz1* does not strongly inhibit invasive growth in the presence of sulfide. However, this initial finding was for one particular experiment, and may reflect experimental variability. Determining whether this result is significant requires statistical analysis.

We adopted a systematic experimental design to identify conditions that maximise the ability to detect sulfide-induced invasive growth, and to assess potential interactions with gene-deletion mutants. We also varied the type of growth medium, the initial level of ammonium (*i*.*e*. nitrogen), and the duration that colonies were allowed to grow (the time point at which plates are washed). In our experimental design, we used both the presence and degree of invasion measures to assess invasive growth across a sequential series of experimental assays, comprising more than 3,000 individual colony images. Statistical analysis of the degree of invasion showed that sulfide significantly increased invasion in the parent AWRI 796 strain, with effects detectable at the 95% confidence level. In addition, we explored the influence of nutrient availability, growth medium, plate washing time, and a selection of gene-deletion mutants. Most of the tested mutants reduced invasion relative to the parent strain. Interaction effects between sulfide and genetic background were also examined. However, we found no evidence that these interactions suppress or enhance the invasive response to sulfide in the majority of mutations. This might indicate the effect of sulfide occurs via a novel pathway. Our results also demonstrated the enhanced information gained from quantitative image analysis in a washed-plate experimental design, compared with binary assessments of invasion alone.

## Results

To validate our measures of invasion, we first analysed images of the parent AWRI 796 strain to examine the effect of ammonium availability (Figure 2)^28^. All colonies displayed evidence of invasion, yielding a value of one for the presence of invasion across all experimental conditions. As ammonium levels decreased, depriving colonies of a key nutrient, we observed an increase in the degree of invasion. This result validated our experimental and analytical approach, as enhanced invasive growth under low-ammonium conditions is well documented^9^. At the same time, it highlighted a limitation of the presence of invasion measure, which cannot distinguish between conditions when invasion occurs in all colonies.

**Figure 2.**
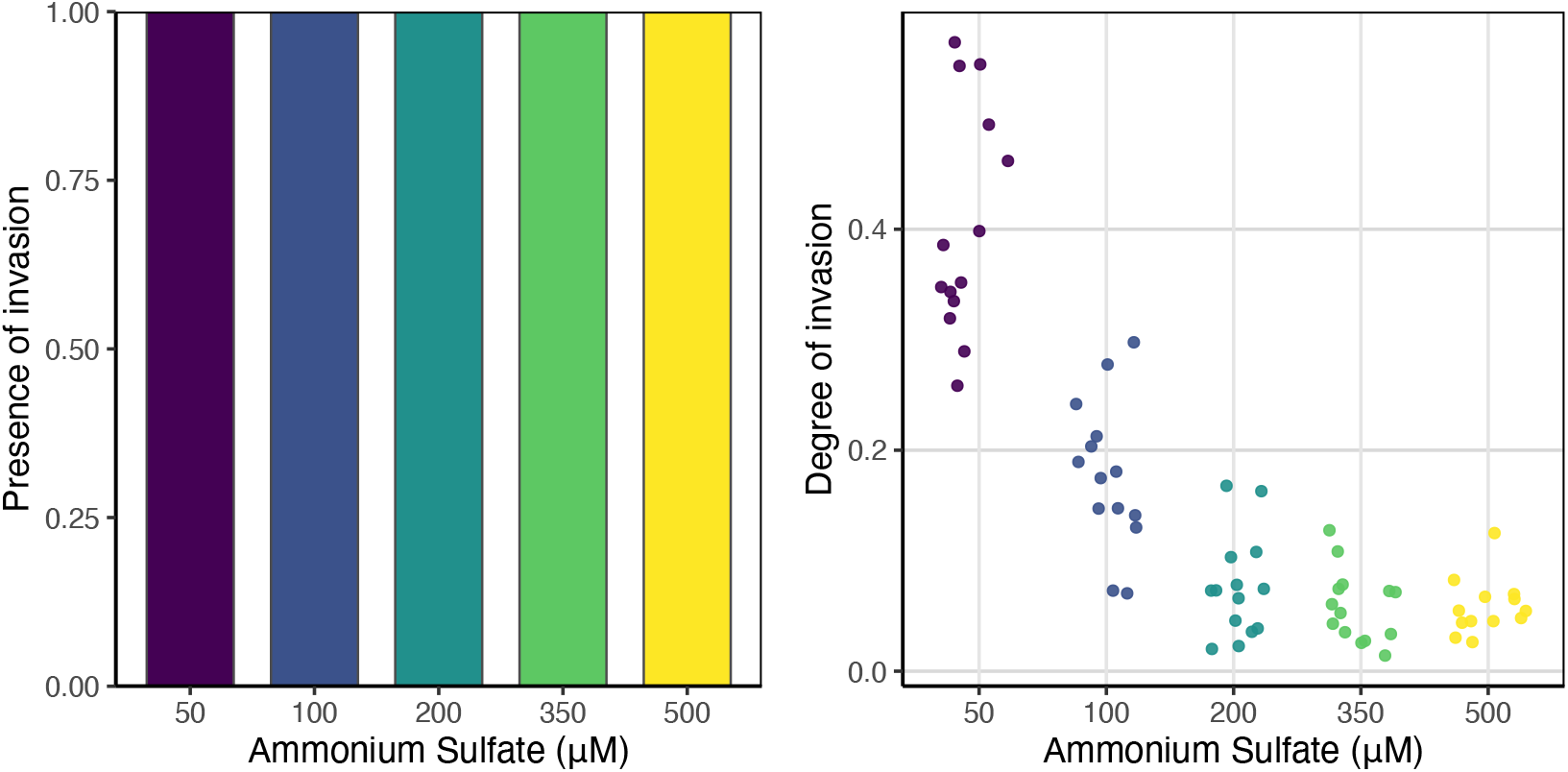
Effect of ammonium sulfate concentrations (50 µM to 500 µM) on invasive behaviour of AWRI 796, 50 µM (purple), 100 µM (blue), 200 µM (dark green), 350 µM (light green), 500 µM (yellow). Left panel: Presence of invasion. Right panel: Degree of invasion. All colonies were grown on 1×SLAD BD agar and washed at day 10 of the assay.

Having established the validity of our measures, we turned to an exploratory analysis of invasive growth across experimental conditions, guided by the patterns observed in Figure 1. Visual inspection revealed general trends in how sulfide exposure, genotype, and growth conditions affect invasive growth in *S. cerevisiae*. We investigated these trends further using statistical analysis, first considering the influence of sulfide on invasive growth in the parent AWRI 796 strain.

### Impact of sulfide on invasive growth of AWRI 796

The first set of experiments was designed not simply to compare conditions, but to systematically identify those under which sulfide-induced invasive growth can be most clearly detected. To achieve this, we varied key experimental factors—including agar type, nutrient availability, incubation time, and medium composition—within a structured factorial design. This approach allowed us to assess not only the direct effects of these factors, but also how they influence the ability to resolve the effect of sulfide on invasive growth. In particular, conditions that produced uniformly high or low levels of invasion (such as Oxoid agar) were less informative for detecting sulfide effects, highlighting the importance of experimental design in establishing conditions where meaningful differences can be observed.

The experiments examined the impact of sulfide concentration on the invasive growth of the parent AWRI 796 strain using a fully factorial design. The variables in these experiments included sulfide levels (0, 400, and 750 µM), ammonium concentration (50, 75, and 100 µM), the duration of growth and thus the day of plate washing (day 3 or day 6), the manufacturer of the agar (Becton Dickinson (BD) or Oxoid), and the final concentration of SLAD medium used for growth (1×SLAD or 2×SLAD). Together, these factors defined a total of 72 experimental conditions.

We began by examining the results for the combination of preculture, incubation time, medium composition and agar source. Broad trends were evident in both the degree of invasion and presence of invasion measures for colonies grown on BD agar (Figure 3). Invasive growth was greatest in colonies grown on the 2×SLAD medium for 6 days (fourth column). The least invasion was observed on day 3 (first and second columns). This shorter time period was insufficient to capture invasive growth, regardless of the other conditions, including SLAD, ammonium, and sulfide concentrations.

**Figure 3.**
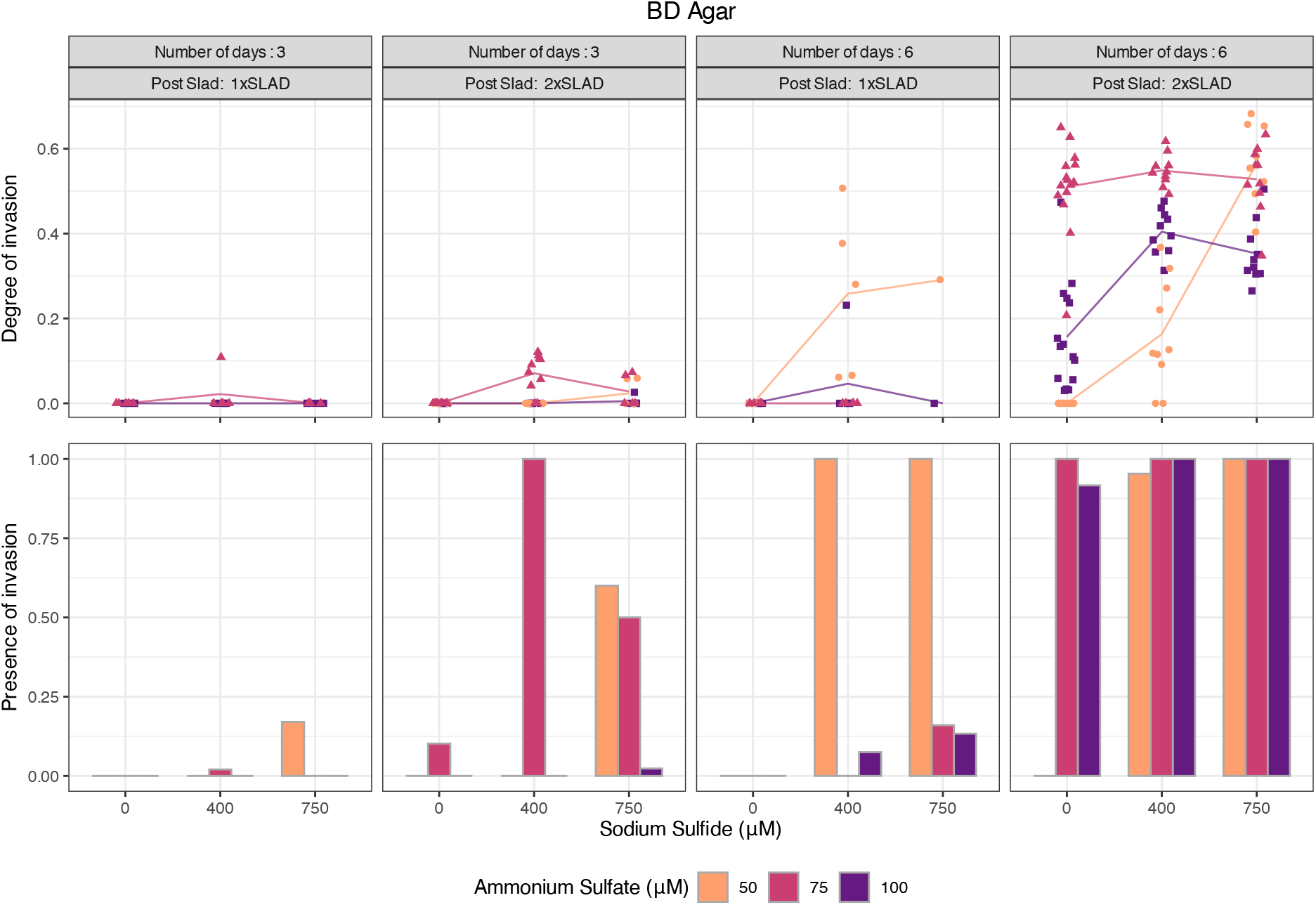
Presence and degree of invasion of AWRI 796 grown on 1 × or 2 × SLAD with Becton Dickinson (BD) agar, with 50 (orange, light grey circles), 75 (mauve, medium grey triangles) or 100 (purple, dark grey squares) µM ammonium sulfate for 3 or 6 days. Line segments connect at the mean value for each sulfide condition.

Before comparing the two invasion measures, it is important to note the difference in the underlying datasets used in their calculation (see Methods). For each experimental condition, the presence of invasion was computed using the total number of colonies on the plate. In contrast, we calculated the degree of invasion for each of a subset of colonies for which image-based measurements were available. For the condition of 50 µM ammonium sulfate, 750 µM sodium sulfide, day 3 plate washing, and 1×SLAD (first column, Figure 3), the presence of invasion was calculated from a total of 47 colonies, whereas the degree of invasion analysis was based on a smaller subset of 5 colonies. This difference in underlying datasets explains an apparent contradiction between the two measures. The markers for the degree of invasion were all zero, while the presence of invasion took a non-zero value of approximately 0.15. Despite this discrepancy, it remained useful to compare both measures across experimental conditions.

Returning to the 2×SLAD condition with plate washing on day 6 (fourth column, Figure 3), the presence of invasion values was high for all ammonium sulfate and sodium sulfide concentrations, except for the 50 µM ammonium sulfate condition in the absence of sodium sulfide. However, the presence of invasion measure provided limited information about the effect of increasing sulfide, whereas the degree of invasion measure revealed clear increases and decreases in the extent of invasion. This limitation of the presence of invasion measure was also evident in Figure 2, where it attained a value of 1 for all ammonium concentrations, thereby masking the decrease in invasion as nutrient levels increased. The degree of invasion measure increased as the sulfide concentration increased for the 50 µM ammonium sulfate condition. Since this change was readily apparent, we considered the day 6 and 50 µM ammonium sulfate experimental conditions to be the best for investigating the effect of sulfide on invasive growth.

In growth medium with Oxoid agar, nearly all experimental conditions produced invasion (second row, Figure 4). In general, the presence of invasion exceeded that observed for the same conditions on BD agar (second row, Figure 3). At first glance, Oxoid medium may appear to be promising for examining the effect of sulfide on invasive growth. However, the trend of increasing degree of invasion (first row, Figure 4) with increasing sulfide concentration was not as strong as that seen for the 2×SLAD condition with plate washing on day 6 and 50 µM ammonium sulfate on BD agar (fourth column, Figure 3). Oxoid agar promoted invasive growth to such an extent that the effect of added sulfide was obscured in these experiments. Hence, BD agar was the more suitable medium for detecting the effect of sulfide on invasive growth.

**Figure 4.**
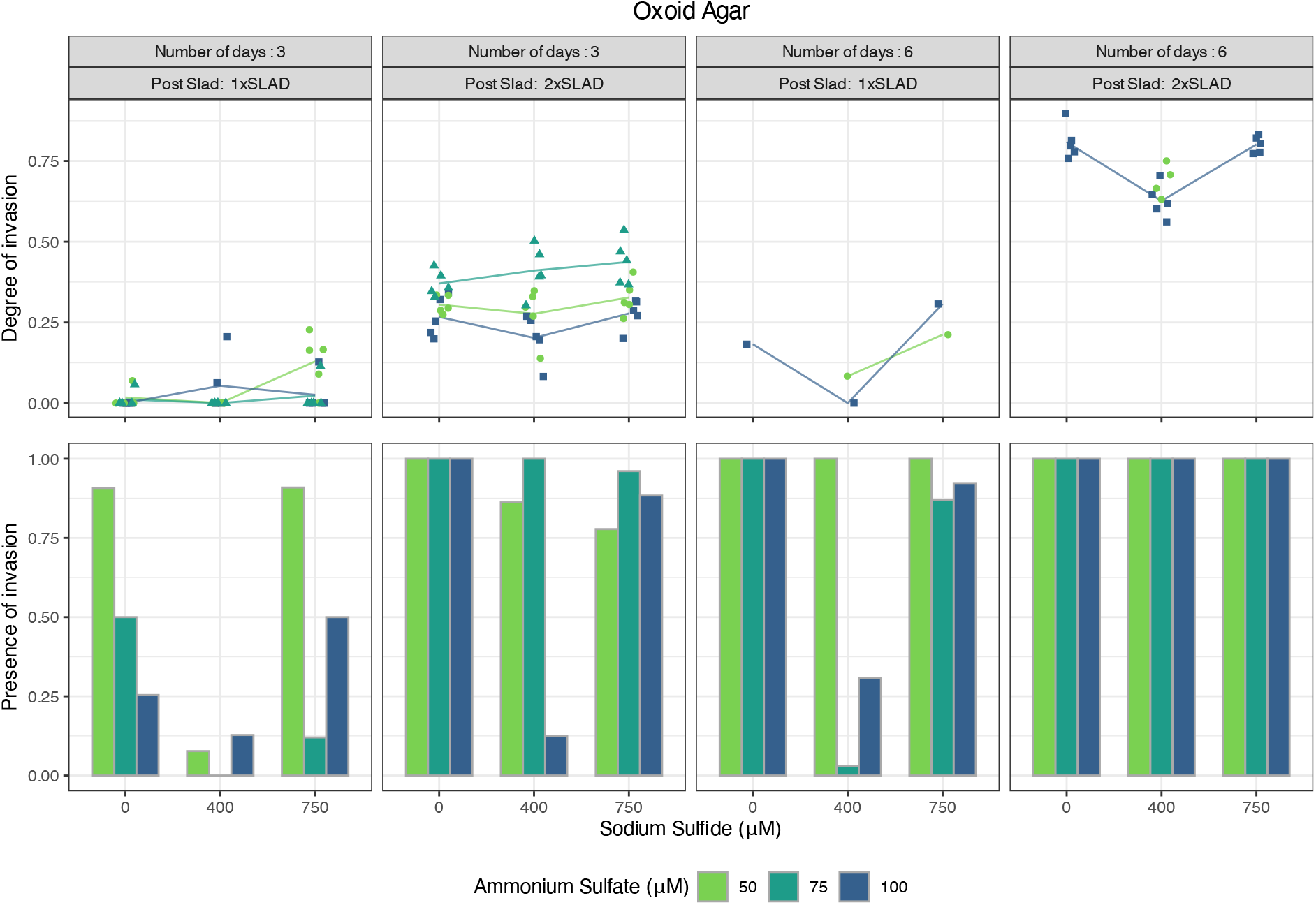
Presence and degree of invasion of AWRI 796 grown on 1 ×or 2 ×SLAD with Oxoid agar, with 50 (green, light grey circles), 75 (teal, medium grey triangles) or 100 (blue, dark grey squares) µM ammonium sulfate for 3 or 6 days. Line segments connect at the mean value for each sulfide condition.

To more rigorously assess the trends in invasion across experimental conditions, we used the degree of invasion measure in this analysis, because it better reflected the overall extent of invasive growth compared to the presence of invasion measure. We performed statistical analysis using a Beta regression (see Methods). A beta regression is appropriate for a rate or proportion outcome^29^, which has values that lie between 0 and 1 (*e*.*g*. the degree of invasion measure). It uses a parametrisation of the Beta distribution that can describe both the precision (spread or variance) of the outcome and the mean of the outcome. In our model, we assumed a constant precision across the experimental conditions, because the small sample size made it difficult to model both precision and mean by different experimental conditions. We instead modelled the mean of the outcome by different experimental conditions, which allowed us to state whether there is a significant difference between conditions with sulfide and conditions without sulfide (and other comparisons between experimental conditions) in terms of the mean degree of invasion.

To assist a reader unfamiliar with the Beta regression to interpret the parameter estimates, we summarised each model through coefficient plots (full parameter estimates are including in tables in the Appendix). The plots summarise the estimated effects of each experimental condition in comparison to a reference condition. In Figure 5, each marker represents the estimated coefficient for a given variable and condition in the model. Horizontal line segments through the markers show the associated 95% confidence intervals, indicating the range of values within which the true parameter is likely to lie. The dashed vertical line at zero denotes where there is no significant difference between the reference category and the category represented on the *y*-axis. Coefficients positioned to the right of this line (positive values) indicate conditions associated with greater invasion relative to the reference condition, whereas those to the left (negative values) indicate reduced invasion. The vertical axis lists the model variables and conditions of the variable that can be interpreted with respect to the reference class. Statistically significant results are shown in yellow (light grey).

**Figure 5.**
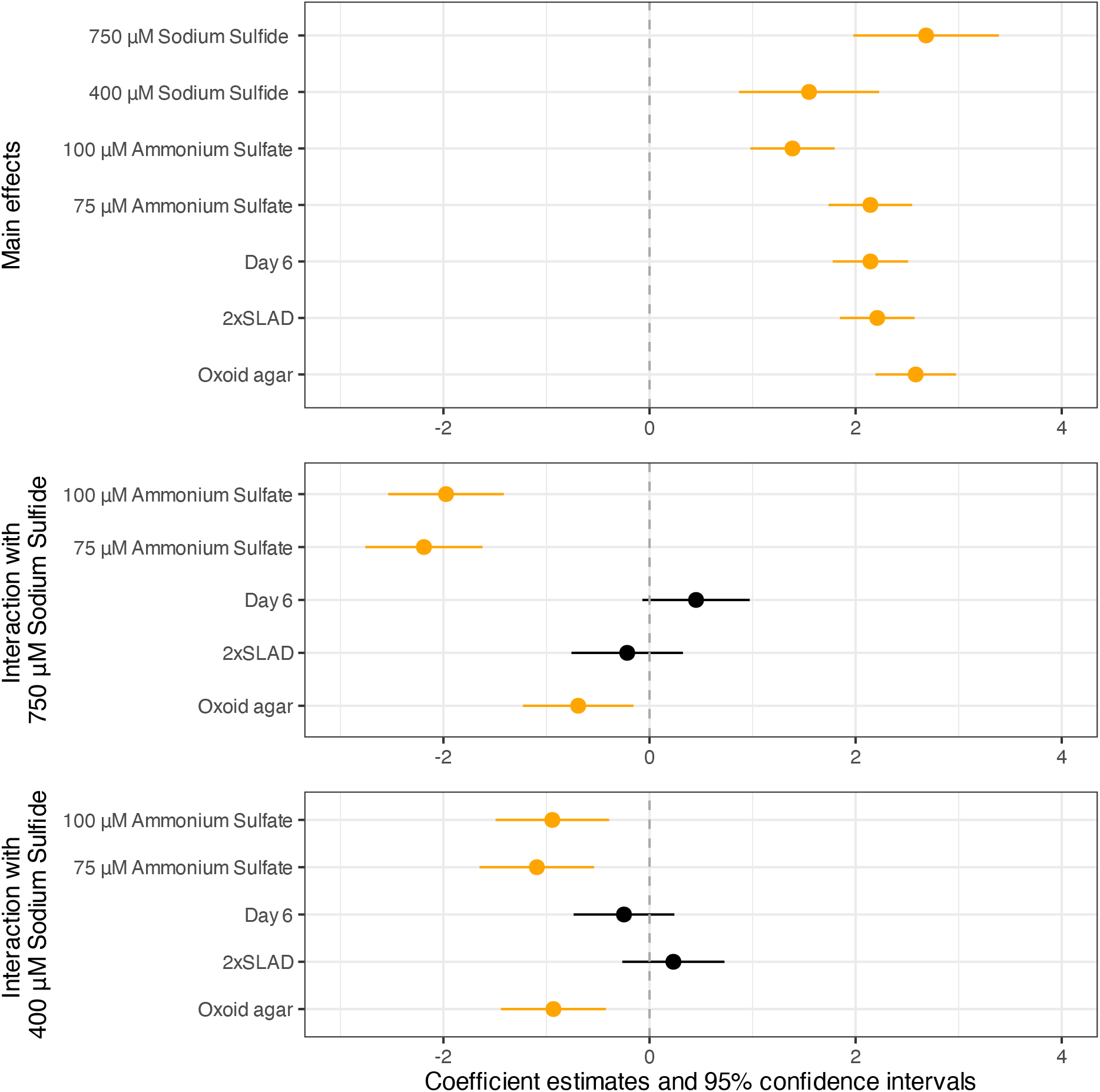
Coefficient plot of the Beta model showing estimated effects of experimental conditions for the results shown in Figures 3 and 4. Each marker represents a coefficient estimate, with horizontal lines indicating 95% confidence intervals. The dashed vertical line at zero marks no effect; positive values indicate increased invasion relative to the reference, negative values indicate decreased invasion. Variables are listed on the vertical axis, and statistically significant effects are highlighted in yellow (light grey). Top panel: Main effects. Middle and bottom panels: Interaction effects with sodium sulfide.

There was a statistically significant increase in invasion for all main effects listed in the top panel of Figure 5. Sulfide, ammonium, day 6 of washing, 2×SLAD, and Oxoid agar all showed higher invasion relative to their respective reference conditions of no sulfide, 50 µM ammonium sulfate, day 3 of washing, 1×SLAD, and BD agar. These results aligned with the observations from Figures 3 and 4. However, the increased invasion for the two ammonium concentrations may seem to contradict the general trend in Figure 2, where invasion decreased with higher ammonium. This discrepancy arose because the range of sulfate values used in the statistical analysis was much smaller than in the visualisation of Figure 2. Indeed, the right panel of Figure 2 showed substantial overlap between the first two conditions, corresponding to the narrower interval between different sulfate conditions used in Beta model analysis. Importantly, we confirmed with statistical significance that adding sulfide increases invasion in the parent AWRI 796 strain.

We next turned to interaction effects between the two sulfide concentrations and the variables listed on the vertical axis in the middle and bottom panels of Figure 5. Both panels showed statistically significant interactions, indicating that sulfide combined with the two higher ammonium concentrations produced significantly less invasive growth relative to the 50 µM ammonium sulfate and zero sulfide reference conditions. However, there was no statistically significant interaction between sulfide and either SLAD concentration or day of washing. The interaction between sulfide and Oxoid agar resulted in significantly less invasion compared with the reference BD agar. This less pronounced relative increase arose because Oxoid agar already promoted substantial invasive growth in the absence of sulfide, leaving less scope for additional enhancement when sulfide was added. This idea is consistent with the raw data, and indicated that 50 µM ammonium sulfate on BD agar was optimal for detecting the effect of sulfide on invasion. These results informed the experimental design used to investigate gene-deletion mutant strains in the next section. We used 2×SLAD and BD agar for all experiments, with two ammonium sulfate concentrations, a single sulfide concentration and multiple plate-washing time points.

### Response of gene-deletion mutants of AWRI 796 to sulfide

In the next set of experiments we aimed to understand the impact of gene deletions on sulfide-enhanced invasive growth. These genes were selected based on their reported involvement in yeast sulfur metabolism. Genes were selected from those known to function in a relevant biochemical pathway or phenotype according to the annotation tools in the *Saccharomyces* Genome Database^26^, and relevant literature mining (Supplementary material, Table 3). These experiments were conducted in three batches. The first batch included the yeast gene deletants *fat1, gup1, nrt1, skp2, soa1* and *yor1*. The second batch included *ccz1, cdh1, cvt16, msa1, pep12, rps8a, tmn3, vps28* and *whi3*, and the third *alr2, ato3, dur3, fui1, mid1, msb2* and *tpo4*. All three batches used the same environment; 2×SLAD with BD agar and experimental manipulations; ammonium sulfate (50 or 75 µM), sodium sulfide concentration (0 or 400 µM), and two plate-washing time points, giving eight experimental conditions for each mutant. The parent AWRI 796 strain was included in all three batches to control for unaccounted laboratory variation. In the first batch, two additional strains, Σ1278b (lab strain) and L2056 (wine strain), were also included for comparison to AWRI 796. In the first batch, the plates were washed on days 3 and 6. In the second and third batches, both parent AWRI 796 strain and gene-deletion mutants were washed on days 4 and 6. All three batches were analysed together in one model. Having established that the count-based presence of invasion measure is limited and not suitable for statistical testing, we focused on the degree of invasion measure and present coefficient plots from a statistical analysis using a Beta regression model (Figure 6).

**Figure 6.**
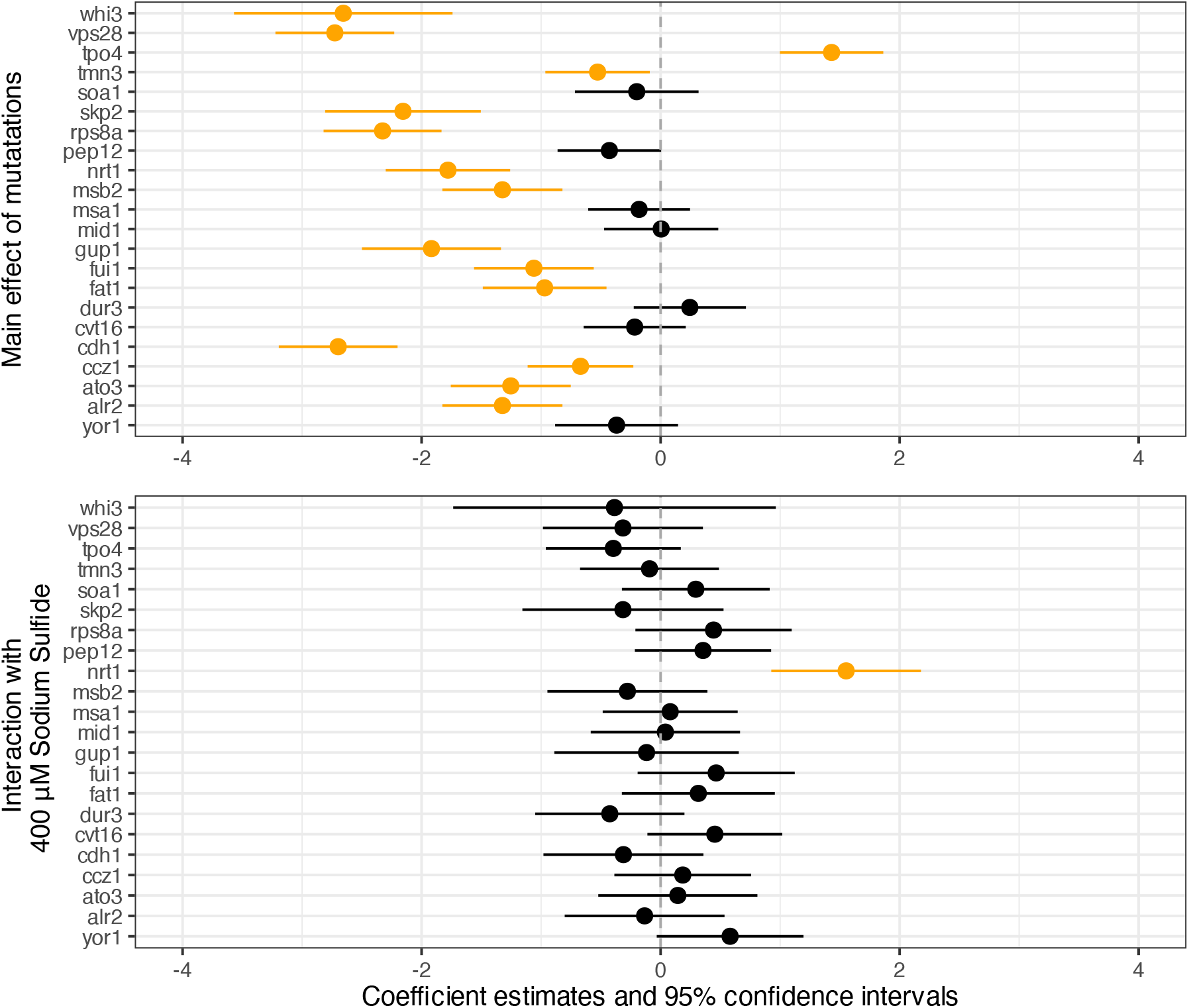
Coefficient plot of the Beta model showing estimated effects for gene-deletion mutants of parent AWRI 796 strain. Each marker represents a coefficient, with horizontal lines indicating 95% confidence intervals. The dashed vertical line at zero marks no effect; positive values indicate increased invasion relative to the reference, negative values indicate decreased invasion. Variables are listed on the vertical axis, and statistically significant effects are highlighted in yellow (light grey). Top panel: Main effects. Bottom panel: Interaction effects with sodium sulfide.

We began by examining the main effects of mutations. Many of the gene deletions led to a significant decrease in invasive growth relative to the parent AWRI 796 strain, with the only exception being the deletion of the *TPO4* gene, which resulted in significantly increased invasion (top panel of Figure 6). This indicated that most of these genes are required for normal invasive growth, while *TPO4* may act as a negative regulator of invasion. In contrast, when considering the interaction between the mutants and 400 µM sodium sulfide, we saw no significant change in invasion relative to the parent strain for most mutants, except for the deletion of *nrt1* (bottom panel of Figure 6). Sulfide at 400 µM did not amplify or reduce invasion in most mutants compared to the invasion response of the parent strain.

To aid interpretation of the interaction effects, we provide a schematic illustration of the no-interaction case, compared with representative positive and negative interactions. No interaction was observed for most mutants, where we could not conclude that the mutant’s response to sulfide differed from that of the parent strain (red and blue lines in Figure 7). The parent strain has a higher baseline level of invasion than the mutants, reflecting the effect of the deleted genes. When sulfide is added, invasion increases in both the parent and mutant strains. Importantly, the increase is of a similar magnitude in each case, which is shown by the parallel (red and blue) lines. In absolute terms, the level of invasion in the mutant with sulfide remains lower than that of the parent with sulfide, reflecting the reduced baseline. In statistical terms, this corresponds to a lack of a significant interaction effect, where the mutations shift the baseline level of invasion, but do not measurably alter the response to sulfide under the conditions tested. For completeness, dashed and dotted lines illustrate positive and negative interaction cases, respectively, where the response to sulfide is either enhanced or diminished in the mutant relative to the parent strain.

**Figure 7.**
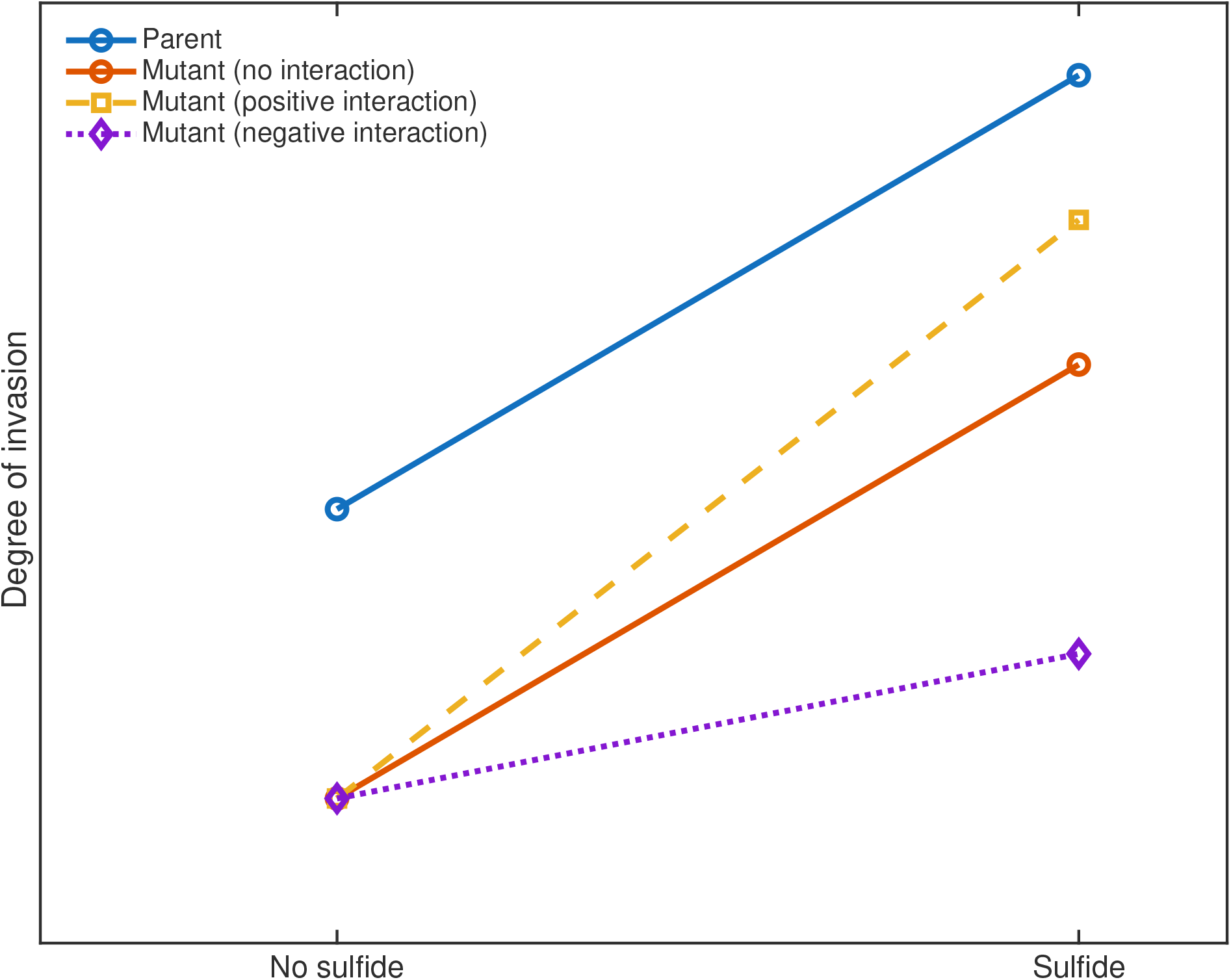
Representative schematic of possible mutant–sulfide interaction effects, where the mutant (lowest marker) exhibits significantly lower invasion than the parent strain (leftmost blue marker).

The lack of statistically significant results in the lower panel of Figure 6 suggests that most mutations studied here did not strongly disrupt the mechanism by which sulfide promotes invasive growth. One possible explanation for this finding is that sulfide acts downstream of the deleted genes in the relevant signalling pathways, so that the effect of sulfide is preserved even when earlier components are removed. Alternatively, sulfide may act through parallel or compensatory pathways that bypass the disrupted genes, allowing invasive growth to be induced via a different route. It is also possible that more subtle effects on the sulfide response exist, but were not detectable given the current sample size and experimental resolution. This interpretation is supported by Figure 1. Without sulfide (S0), the parent strain (a, b) exhibited a dense central colony with clear invasive growth visible after washing, whereas the ccz1 mutant (e, f) showed reduced invasion, consistent with a lower baseline invasive capacity. In the presence of sulfide (S400), both the parent (c, d) and the mutant (g, h) showed increased invasive growth following washing, with more extensive filamentous structures apparent. Importantly, although the mutant colonies remained less invasive than the parent in absolute terms under sulfide conditions, the qualitative increase in invasion upon addition of sulfide was evident in both strains. Thus, Figure 1 visually reinforces the statistical finding that the mutation reduced the overall level of invasion, but did not significantly diminish the observed response to sulfide.

To control for comparisons of gene deletions with the parent AWRI 796 strain, we also tested two additional strains, Σ1278b and L2056 (Figure 8). The Σ1278b strain is well known to readily invade, and as such is commonly used in invasive growth studies. However, no prior data existed regarding its invasive response to the addition of sulfide, so we had no specific expectations for the interaction effect with sulfide. The L2056 strain is a wine strain, like AWRI 796, and its invasive properties had not been previously characterised. Examining these strains also provided a broader picture of invasive behaviour in yeast and the potential interaction effects with sulfide relative to the AWRI 796 strain. As expected, the Σ1278b strain invaded more than the AWRI 796 strain (top panel, Figure 8). Conversely, the L2056 strain showed significantly lower invasive growth, a result warranting further investigation. When examining the interaction effects between the Σ1278b and L2056 strains and sulfide, relative to the AWRI 796 strain, there was no significant difference for Σ1278b, and increased invasion for L2056. These results indicated that the response to sulfide can be strain-specific. The less invasive L2056 strain showed enhanced invasive growth in the presence of sulfide. In contrast, the strain Σ1278b is already highly invasive, and showed little or no additional invasion in the presence of sulfide.

**Figure 8.**
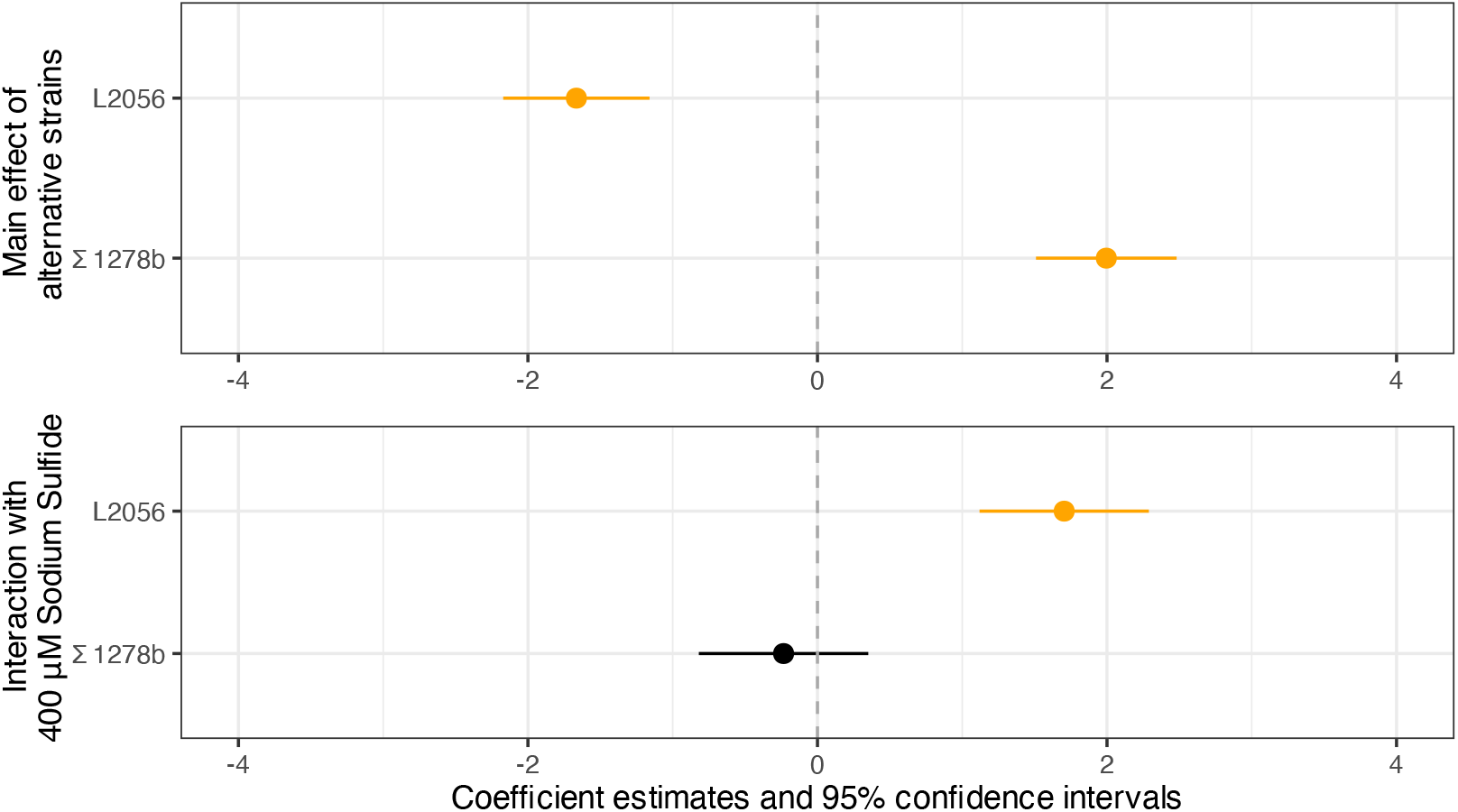
Coefficient plot of the Beta model showing estimated effects for strains Σ1278b and L2056. Each marker represents a coefficient, with horizontal lines indicating 95% confidence intervals. The dashed vertical line at zero marks no effect; positive values indicate increased invasion relative to the reference, negative values indicate decreased invasion. Variables are listed on the vertical axis, and statistically significant effects are highlighted in yellow (light grey). Top panel: Main effects. Bottom panel: Interaction effects with sodium sulfide.

Significant main effects were also found for increased invasive growth in the presence of sulfide (relative to its absence) and for day 4 and day 6 washing (relative to day 3), as illustrated in Figure 9. These results aligned with our previous analysis (see Figure 5). There was no measurable difference between the 75 µM and the reference 50 µM ammonium sulfate concentrations. This similarity was most likely due to the small difference between these two concentrations. There was a significant difference between experimental batches, with the second batch showing greater invasion and the third batch less invasion than the first. This suggested batch-specific effects, such as subtle variations in experimental conditions, media preparation, or culture handling. This possible variability highlighted the importance of controlling with the parent AWRI 796 strain in every experiment to account for batch to batch variability in the analysis.

**Figure 9.**
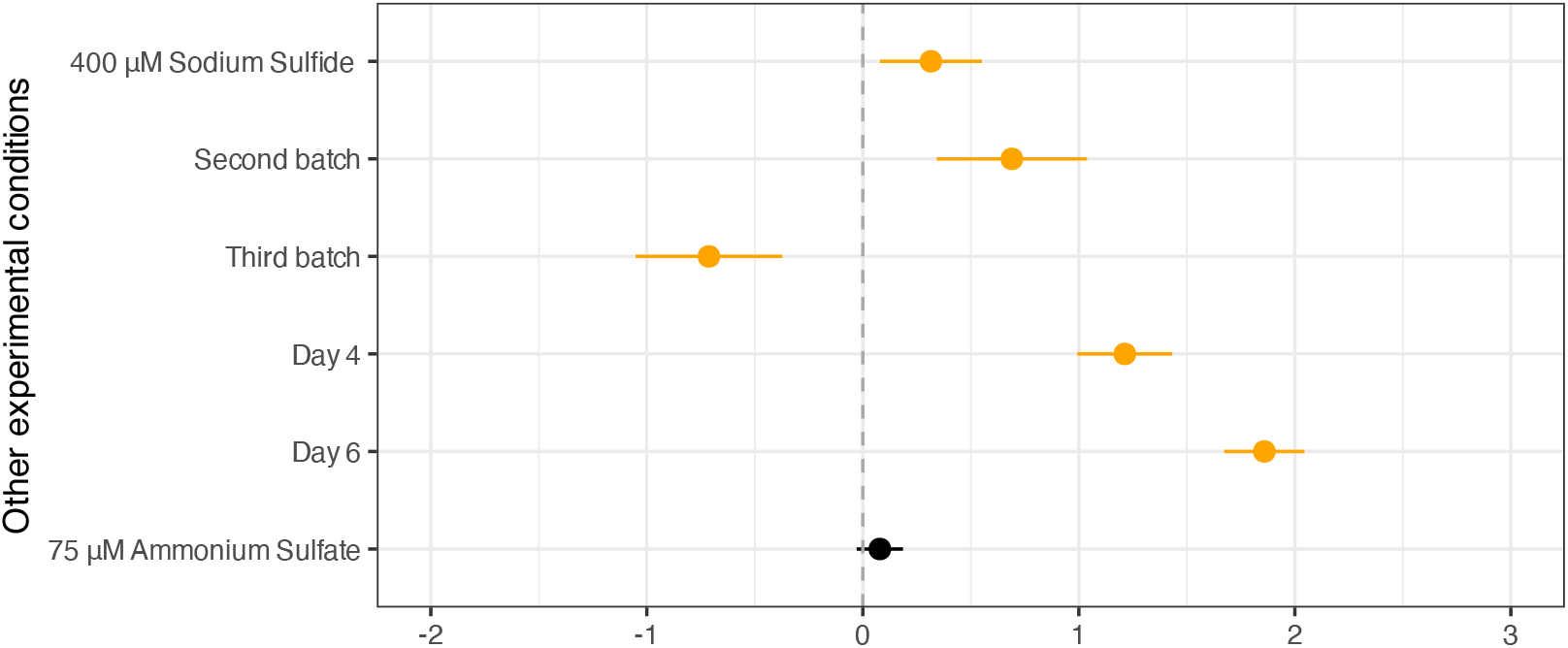
Coefficient plot of the Beta model showing estimated main effects for the experimental conditions in the first, second and third batches of experiments in this section of the Results. Each marker represents a coefficient, with horizontal lines indicating 95% confidence intervals. The dashed vertical line at zero marks no effect; positive values indicate increased invasion relative to the reference, negative values indicate decreased invasion. Variables are listed on the vertical axis, and statistically significant effects are highlighted in yellow (light grey).

### Impact of pre-SLAD on invasive growth in AWRI 796

We examined the effect of pre-culture conditions on invasive growth in the parent AWRI 796 strain before examining the gene-deletion mutants. In all previous experiments, cells were preconditioned under nitrogen-limiting conditions using a pre-1×SLAD medium prior to plating on assay plates (see Methods). We next compare this with a pre-2×SLAD, to assess the effect of pre-culture conditions on the invasive growth in the parent AWRI 796 strain. Once plated, assays were performed on 2×SLAD prepared with BD agar as the assay medium. Alongside the two pre-SLAD conditions (pre-1×SLAD and pre-2×SLAD), we also investigated two ammonium concentrations (50 µM and 75 µM), a single non-zero sulfide concentration (400 µM), and the day of plate washing. Plates from the pre-1×SLAD condition were washed on days 3, 4, and 6, whereas those from the pre-2×SLAD condition were washed on days 3 and 6. The results were summarised in the coefficient plot (Figure 10) from a Beta regression model (see Methods).

**Figure 10.**
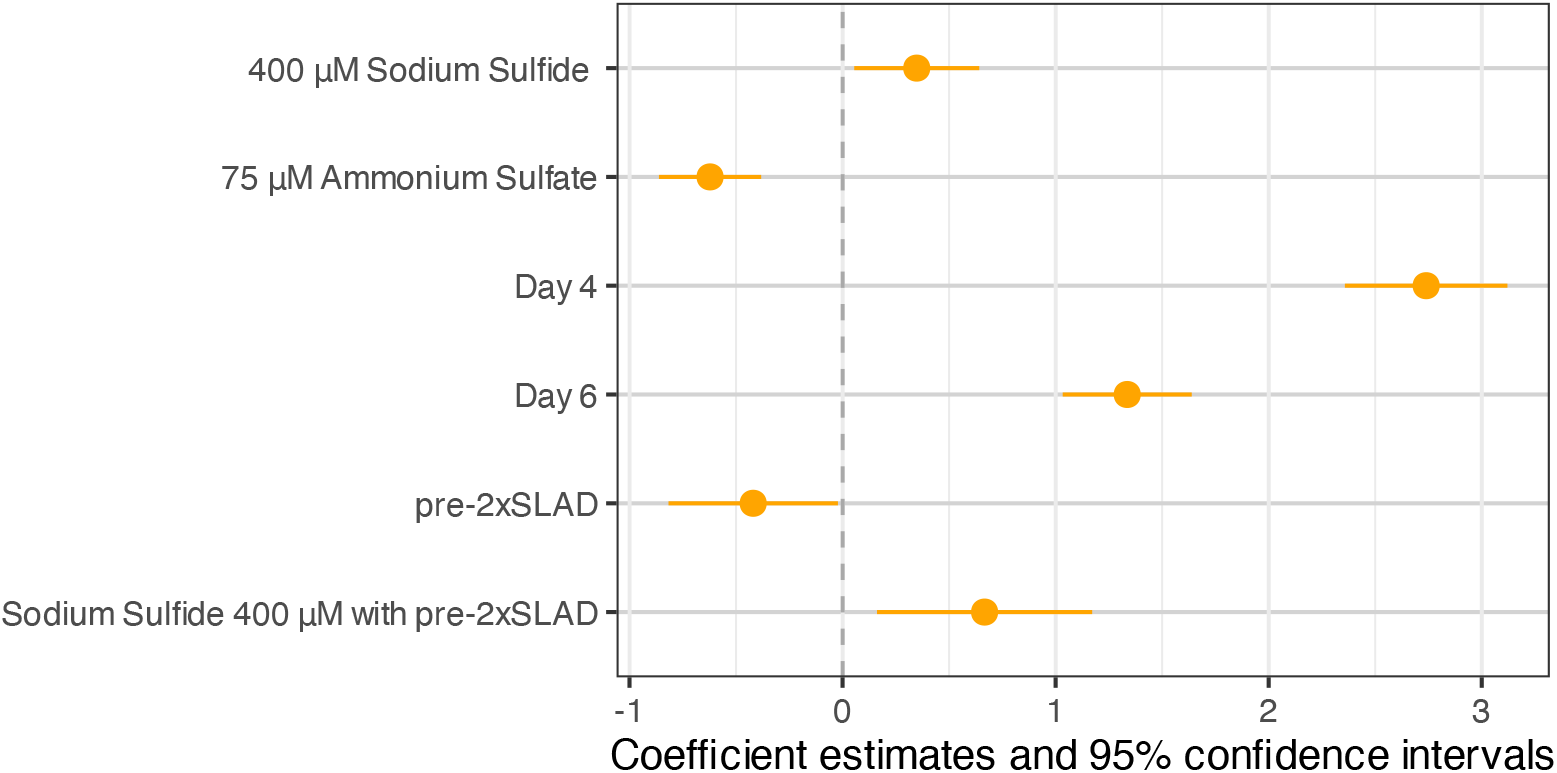
Coefficient plot of the Beta model showing estimated effects for the pre-culture conditions, pre-1×SLAD and pre-2×SLAD, in the parent AWRI 796 strain. Each marker represents a coefficient, with horizontal lines indicating 95% confidence intervals. The dashed vertical line at zero marks no effect; positive values indicate increased invasion relative to the reference, negative values indicate decreased invasion. Variables are listed on the vertical axis, and statistically significant effects are highlighted in yellow (light grey). Top four markers: Main effects. Bottom marker: Interaction effects with sodium sulfide.

The pre-2×SLAD condition showed significantly lower invasive growth as a main effect compared with the reference pre-1×SLAD condition (second from bottom marker, Figure 10). However, there was also significantly greater invasive growth in the interaction between sulfide and the pre-2×SLAD condition (bottom marker, Figure 10). This suggests a pre-2×SLAD provided a more sensitive background for detecting sulfide-induced invasive growth, despite its lower main effect on invasion. One possible explanation for this result is that stronger nitrogen limitation during preconditioning alters the regulation of genes associated with invasive growth, such as *FLO11*^30^. This may change how cells respond to sulfide and allow a larger increase in invasion. For the remaining main effects, we observed results consistent with the analysis so far. As expected, the presence of sulfide (reference, no sulfide) and washing on days 4 and 6 (reference, day 3) were both associated with significantly greater invasive growth.

### Impact of pre-2×SLAD and on invasive growth in gene-deletion mutants of AWRI 796

Given the enhanced invasive growth observed for the interaction between sulfide and the pre-2×SLAD condition (bottom marker, Figure 10), we next examined how pre-culturing in pre-2×SLAD affects the invasive behaviour of gene-deletion mutants derived from the parent AWRI 796 strain (Supplementary material, Table 3). In this set of experiments, BD agar with 2×SLAD was used as the growth medium, together with three ammonium concentrations (0, 50 and 75 µM), a two sulfide concentrations (0 and 400 µM), with plate washing performed on days 4 and 6 in a fully factorial design.

We investigated the response of gene-deletion mutants derived from the parent AWRI 796 strain (*alr2, dur3, nrt1, tmn3, pma2, put4, sac3*), with results shown in the Beta model coefficient plots of Figure 11. All but one mutant, *dur3*, showed significantly less invasion than the reference parent AWRI 796 strain. For comparison, four mutants (*alr2, dur3, nrt1, tmn3*) were also preconditioned with pre-1×SLAD (see Figure 6), and we obtained similar results under the pre-2×SLAD condition (middle panel Figure 11).

**Figure 11.**
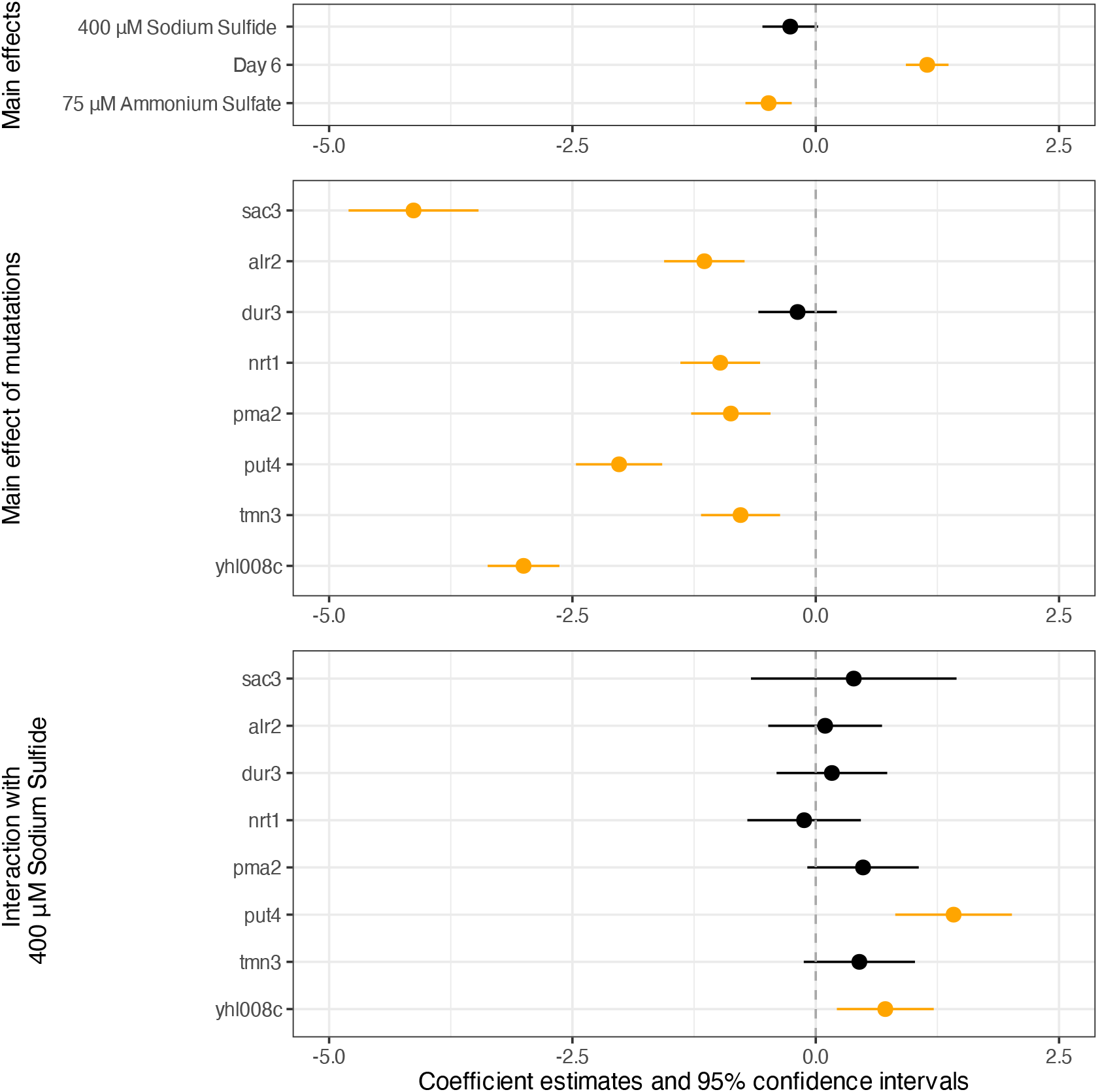
Coefficient plot of the Beta model showing estimated effects for the pre-culture conditions, pre-2 ×SLAD and BD agar with 2 ×SLAD growing medium, for gene-deletion mutants of parent AWRI 796 strain. Each marker represents a coefficient, with horizontal lines indicating 95% confidence intervals. The dashed vertical line at zero marks no effect; positive values indicate increased invasion relative to the reference, negative values indicate decreased invasion. Variables are listed on the vertical axis, and statistically significant effects are highlighted in yellow (light grey). Top panel: Main effects, sulfide, day of washing, sulfate. Middle panel: Main effects, mutants. Bottom panel: Interaction effects of mutants with sodium sulfide.

Finally, we examined the main effects of 400 µM sodium sulfide (interpreted with reference to no sulfide), day 6 plate washing (reference, day 4), and 75 µM ammonium sulfate (reference: 50 µM ammonium sulfate), as shown in the top panel of Figure 11. Significant effects were observed for the washing day and ammonium sulfate concentration, consistent with our previous findings. However, unlike earlier experiments, the main effect of sulfide was not significant. We did observe significant interaction effects between sulfide and the gene-deletion mutants *put4* and *yhl008c* in this set of experiments. Thus, under the conditions tested here, sulfide did not measurably influence invasive growth overall, but it did increase invasive growth for specific mutants. The absence of a main effect may reflect the pre-SLAD treatment or general increased experimental or biological variability in this set of experiments. Since this pattern was observed only in this final set of experiments, the result should be interpreted with caution.

## Discussion

We conducted a systematic experimental analysis of the factors influencing invasive growth in the wine yeast strain AWRI 796, with particular emphasis on the role of sulfide, genetic makeup, and environmental conditions. A principal outcome of this study is the identification of experimental conditions under which sulfide-induced invasive growth can be reliably detected and quantified. Using our experimental design, we confirmed that sulfide is an environmental signal that enhances invasive growth in AWRI 796 under nitrogen-limiting conditions. This effect was observed consistently across all experimental conditions except for the final set of experiments involving pre-2×SLAD preconditioning. In that case, we suspected that relatively low sample numbers and general variability across colonies resulted in a type 2 error.

Analysis of gene-deletion mutants derived from the parent AWRI 796 strain revealed that genetic factors exert a dominant influence on invasive behaviour. Most mutants exhibited significantly reduced invasion relative to the parent strain (Figures 6,11) indicating that the deleted genes contribute positively to the invasive response. Importantly, interaction analyses provided little statistical evidence that the studied mutations changed the overall response to sulfide. This suggested that these deletions largely determine invasive capacity overall, with no statistically significant difference between how the mutants and the parent respond to sulfide. In other words, sulfide modulated invasion in the parent strain and mutant strains similarly, despite general differences in level of invasion.

Comparisons with additional strains further demonstrated the importance of genetic background (Figure 8). Strain Σ1278b,which is well known for strong filamentous and invasive growth, exhibited higher invasion than AWRI 796. This result was consistent with established expectations and validated our experimental framework. By contrast, the wine strain L2056 displayed significantly lower invasion, highlighting substantial diversity in invasive capacity among wine strains. Interestingly,sulfide did not further enhance invasion in Σ1278b, but increased invasion in L2056, suggesting that sulfide responsiveness itself varies across yeast strain. These results indicated that both intrinsic invasive capacity and responsiveness to environmental cues are strain dependent.

Environmental and methodological conditions also played an important role. Incubation time before washing, nutrient composition, agar type, and experimental batch all influenced invasion, emphasising the sensitivity of this phenotype to growth conditions. As expected, the timing of plate washing consistently affected the measured degree of invasion, likely reflecting differences in colony maturation and size at different time points. The absence of a measurable difference between similar ammonium concentrations suggested that relatively small changes in nutrient levels may not noticeably alter invasive behaviour (Figure 5). This was consistent with the experimental results presented in the Introduction, where larger differences in nutrient concentration produced the expected decrease in invasion as nutrient availability increased but smaller differences were much less noticeable.

Pre-culture conditions prior to plating were also important determinants of sulfide responsiveness. Preconditioning in pre-2×SLAD resulted in lower invasive growth as a main effect, but produced a stronger enhancement when sulfide was present. Therefore, this condition provided a more sensitive background for detecting sulfide-induced invasion. One possible explanation for this effect is that stricter nitrogen limitation during pre-culture primes cells for environmental signalling upon transfer to assay plates. This finding highlighted the importance of physiological state prior to plating, which may influence the activation of pathways associated with filamentous growth and invasion.

The variability observed across strains and conditions suggested that invasive growth is a highly variable phenotype shaped by both genetic background and environmental cues. We provided substantive evidence that sulfide increases the invasive growth in most experimental conditions, although measuring this invasive growth was difficult. We also showed that while a number of mutants inhibit overall invasive growth, we did not find a mutant that inhibited the impact of sulfide. Further replication and increased samples sizes are needed to confirm theses findings.

Understanding how sulfide influences invasive behaviour has potential implications for industrial fermentation environments, where sulfide production and nitrogen limitation commonly occur. The differential responses observed among wine yeast strains indicate that both strain selection and environmental management could influence yeast behaviour during fermentation. Future work should therefore aim to identify the molecular mechanisms underlying sulfide sensing and its interaction with pathways controlling adhesion and filamentation, and aim to clarify how these processes vary across genetic backgrounds and environmental conditions. Substantial within-condition variability highlights the need for increased replication in future experiments, and suggests that our results must be interpreted cautiously. Future studies could build on this experimental design framework by increasing sample sizes and leveraging quantitative modelling approaches to better capture variability and further refine our understanding of invasive growth.

## Methods

We used a broad exploratory approach to examine the factors that influence invasive growth in yeast, with particular emphasis on the role of sulfide. The goal was to identify how the presence of sulfide influences invasive growth with varying nutrient availability, genetic background, and pre-culture conditions. We aimed to provide a more comprehensive understanding of the mechanisms governing this invasive phenotype. In this section, we describe the experimental procedures, image processing methods, quantification of invasive growth, and the statistical analyses used in the study.

### Experimental methods

The relevant experimental details included the yeast strains and media employed, the cultivation procedures for invasive growth assays, and the microscopy-based approaches used to image surface and invasive growth.

#### Strains and Media

The prototrophic diploid *Saccharomyces cerevisiae* commercial wine strains AWRI 796 (Maurivin) and Lalvin Rhône 2056^®^ (L2056, Lallemand) and the laboratory strain, Σ1278b (prototrophic diploid) were used in these experiments. Commercial strains, as active dry wine yeast preparations were rehydrated in 100 mL of YPD medium (10 g L^−1^ yeast extract, 20 g L^−1^ bacteriological peptone and 20 g L^−1^ glucose) and incubated overnight at 28 °C. Yeast cultures were stored at −80 °C in the presence of 26 % glycerol.

Synthetic Low Ammonium Dextrose (SLAD) liquid and agar media were prepared as per^15^, and used at either 1×SLAD or 2×SLAD as described, with ammonium sulfate concentrations of 50, 75 or 100 µM and with sodium sulfide (Sigma Aldrich, Cat No 13468) at 0, 400 or 750 µM. SLAD was solidified with the addition of 2 % agar (BD Bacto Agar, Becton Dickinson, Cat No. 233520), unless specified as Oxoid™ bacteriological Agar (Oxoid, Cat No LP0011T). Solidified SLAD media was prepared within 24 hours of use.

Gene-deletion mutants in AWRI 796 were constructed essentially as described in^31^ with the following modifications. Individually *ALR2, ATO3, CCZ1, CDH1, CVT16, DUR3, FAT1, FUI1, GUP1, MID1, MSA1, MSB2, NRT1, PEP12, PMA2, PUT4, RPS8a, SAC3, SKP2, SOA1, TMN3, TPO4, VSP28, WHI3, YHL008c* and *YOR1* were disrupted in AWRI 796. Since AWRI 796 is diploid, transformants resistant to geneticin (0.2 g L^−1^) were sporulated and asci dissected. The target gene alleles (disrupted or wildtype) were then determined from cultures derived from each spore of the four spore asci by Polymerase Chain Reaction with gene specific primers pairs, as described by the yeast deletion project^32^. Strains where the target gene was disrupted from both DNA strands were selected for further studies. For simplicity, yeast gene-deletion mutants are notated in the form *alr2* rather than the conventional, AWRI 796 *alr2△::KanMX4*.

#### Microbial cultivation for invasive growth assays

Yeast were cultured from glycerol stocks in 2 mL of 1×SLAD liquid medium for two overnights at 28 °C with agitation (120 rpm). Cells were subsequently inoculated into 5 mL of fresh 1×SLAD (or where specified 2×SLAD liquid medium) at 1 × 10^5^ cells*/*mL and incubated for 18 hours at 28 °C with agitation (120 rpm) to obtain an exponential phase culture. Dilutions, in phosphate buffered saline (PBS), calculated to contain between 50 to 100 cells, were then spread on 2×SLAD agar (or 1×SLAD agar where indicated) and the plates incubated at 28 °C for the times specified (typically 3 and 6 days).

#### Recording of surface and invasive colony growth with microscopy

Yeast colonies or cultures were observed at 40× magnification using a Nikon Eclipse 50i microscope and imaged using a Digital Sight DS-2MBWc camera and NIS-Elements F 3.0 imaging software (Nikon) at the time points stated. At this time, manual counting of total colonies and the number with detectable invasive growth from each plate was recorded. This was performed by the investigator looking for dark patches (invasive growth) against lighter surface growth, as observed in bright-field microscopy. Typically, plates had 30-50 colonies per plate. Up to 15 colonies per plate were then chosen at random, numbered and their position marked. We ensured those selected were well-separated from neighbouring colonies to decrease the potential influence of neighbouring colonies upon colony morphology. All colonies were then washed using a stream of ultrapure water, thus removing the cells from the surface, and enabling subsequent imaging of the invasive growth of the selected colonies into the agar medium. The total amount of invasively growing colonies was also manually confirmed after plate washing, this was again aided by bright-field microscopy.

### Image processing

We took two grayscale TIFF images of each colony, one before and one after washing. These images were processed using our custom-built, open-source software, TAMMiCol^16^, to convert colony images into binary format prior to further processing, as described below. Pre-wash images were processed using the TAMMiCol *connected* method setting, while post-wash images were processed using the *unconnected* setting. In some cases, adjustments to the threshold parameter were necessary to compensate for variations in image contrast across the TIFF files. Post-wash images contained small, low-contrast background artefacts that were not clearly distinguishable from the surrounding agar (Figure 12(a)), and these artefacts were not always removed during processing with TAMMiCol (Figure 12(b)). Therefore, additional processing was necessary to remove these artefacts (connected components with 10,000 pixels or fewer) using the Matlab function `bwareaopen()` (Figure 12(c)). Additional examples of the three-stage image processing are shown in Figure 13.

**Figure 12.**
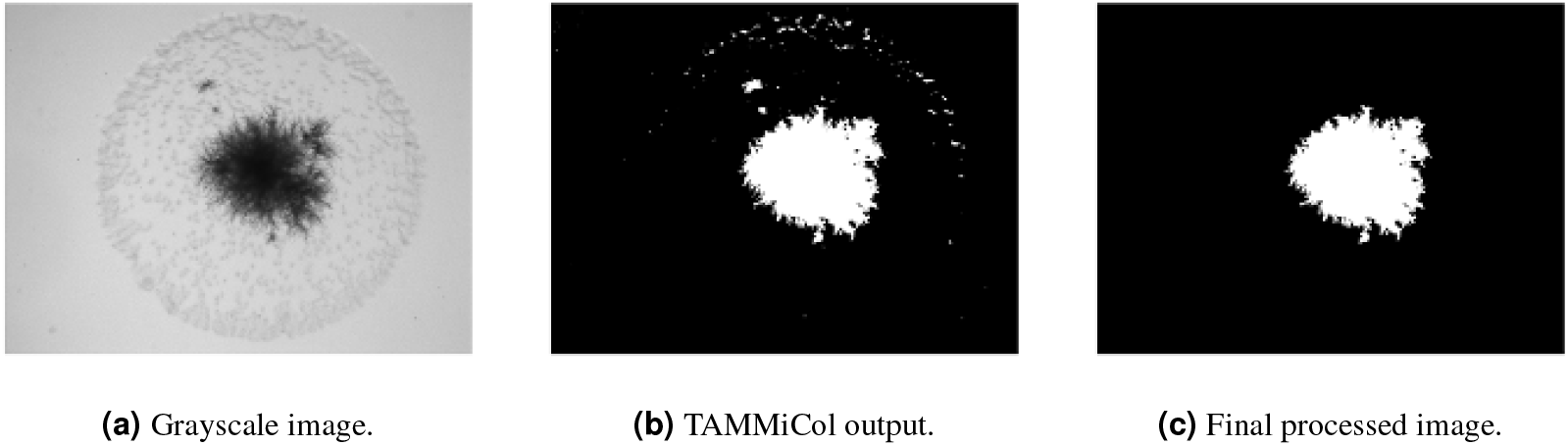
The three-step process of processing and cleaning experimental grayscale images for spatial analysis.

**Figure 13.**
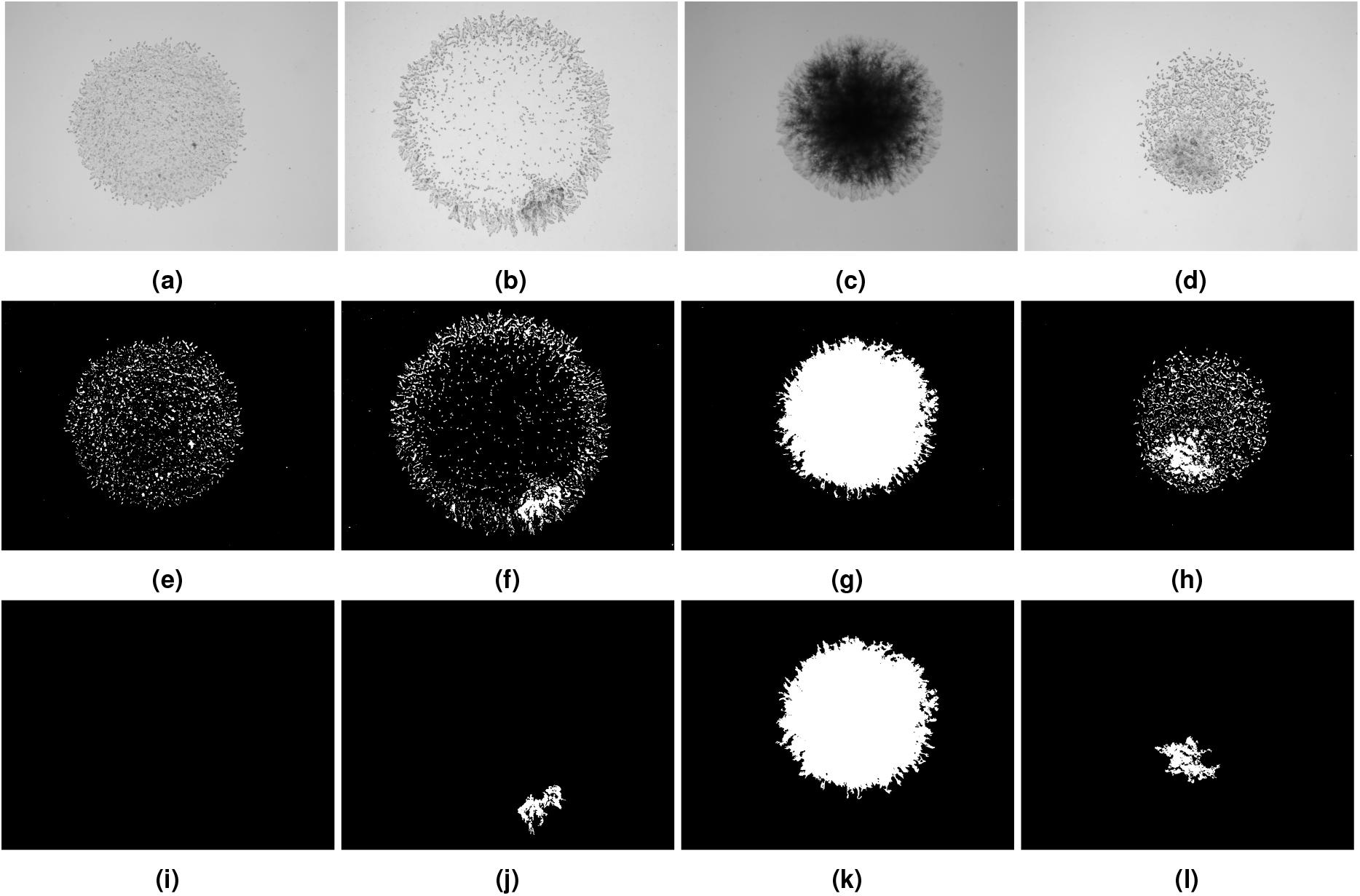
Four representative examples (columns) illustrating the three-stage image processing workflow for post-washed images. Row 1: Grayscale TIFF images. Row 2: Binary images obtained following TAMMiCol processing. Row 3: Final processed images after removal of small artefacts using `bwareaopen()`.

### Measurement of invasion

We used two measurement techniques to quantify invasive growth in our experimental design. The first was a human-based assessment that captured the presence of invasion, in which the total number of colonies, *N*_*j*_, on the *j*th plate was counted prior to washing. Following washing, plates were visually inspected for *any* evidence of invasive growth, and the number of invasive colonies, *n*_*j*_, was recorded. The presence of invasion was then defined as

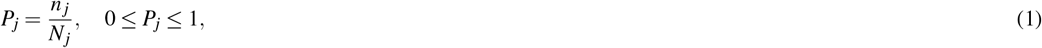

which represents the proportion of colonies exhibiting invasive growth on the *j*th plate (or *j*th experimental condition).

The degree of invasion measure was based on a subset of colonies, *M*_*j*_ *≤ N*_*j*_, on the *j*th plate. Colonies were selected for analysis only if they were sufficiently spaced from neighbouring colonies to allow accurate image processing. Pre- and post-washed images were processed as described in the previous section on image processing. Let the pre- and post-washed areas of the *i*th colony in the subset *M*_*j*_ on the *j*th plate be denoted by *A*_*i, j*_ and *a*_*i, j*_, respectively. The degree of invasion for the *i*th colony was then defined as

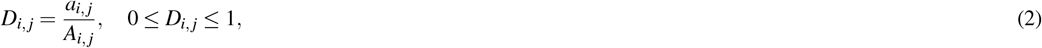

which represents the ratio of the washed area to unwashed area, for each colony (in the sample) on the plate.

The count-based presence of invasion measure yielded a single value for each plate under a given condition, whereas the area-based degree of invasion produced a set of values corresponding to individual colonies within the sampled subset on each plate. As discussed in the Results, the degree of invasion measure was more amenable to statistical analysis.

### Experimental manipulations

The experiments were completed with 5 different cohorts or batches, plus the original nutrient study shown in the introduction. Since we observed some overall differences between the batches, we summarise each batch and its manipulations here. The first experimental batch is presented in Section “Impact of sulfide on invasive growth of AWRI 796”. It consisted of only the parent AWRI 796 strain, with a (3 × 3 × 2 × 2 × 2 = 72 fully factorial design varying the sodium sulfide levels (0, 400, and 750 µM), ammonium sulfate concentration (50, 75, and 100 µM), the duration of growth and thus the day of plate washing (day 3 or day 6), the manufacturer of the agar (Becton Dickinson (BD) or Oxoid), and the final concentration of SLAD medium used for growth (1×SLAD or 2×SLAD). A pre-1×SLAD was used for all conditions. On average there were 45 colonies per plate before washing and 25 colonies that were growing invasively. An average of 6 colonies were imaged in each condition.

The second experimental batch is presented in Section “Response of gene-deletion mutants of AWRI 796 to sulfide”. It consisted of the AWRI 796 strain, Σ1278b and L2056 strains and yeast gene deletants of the AWRI 796 strain *fat1, gup1, nrt1, skp2, soa1* and *yor1*. The experimental design for each strain/mutant was a (2 × 2 × 2 = 8) fully factorial design varying ammonium sulfate (50 or 75 µM), sodium sulfide (0 µM or 400 µM) and plates washed on day (3 or 6). All conditions used pre-1×SLAD with a 2×SLAD BD agar. On average, there were 32 colonies per plate before washing and 18 colonies that were growing invasively. An average of 5 colonies were imaged in each condition for each mutant/strain.

The third experimental batch is presented in Section “Response of gene-deletion mutants of AWRI 796 to sulfide”. It consisted of the AWRI 796 strain and yeast gene deletants of the AWRI 796 strain *ccz1, cdh1, cvt1, msa1, pep1, rps8, tmn3, vps2* and *whi3*. The experimental design for each strain/mutant was a (2 × 2 × 2 = 8) fully factorial design varying ammonium sulfate (50 or 75 µM), sodium sulfide (0 and 400 µM) and plates washed on day (4 or 6). All conditions used pre-1×SLAD with a 2×SLAD BD agar. On average there were 37 colonies per plate before washing and 24 colonies that were growing invasively. An average of 5 colonies were imaged in each condition for each mutant.

The fourth experimental batch is presented in Section “Response of gene-deletion mutants of AWRI 796 to sulfide” and “Impact of pre-SLAD on invasive growth in AWRI 796”. It consisted of the AWRI 796 strain and yeast gene deletants: *alr2, ato3, dur3, fui1, mid1, msb2* and *tpo4*. The experimental design for each strain/mutant was a (2 × 2 × 2 = 8) fully factorial design varying ammonium sulfate (50 or 75 µM), sodium sulfide (0 or 400 µM) and plates washed on day (4 or 6). A small number of plates (4) of the parent colonies were washed on day 3. For the parent strain only, an additional manipulation was added using a pre-2×SLAD to compare against the standard pre-1×SLAD. All other conditions used pre-1×SLAD with a 2×SLAD BD agar. On average there were 37 colonies per plate before washing and 19 colonies that were growing invasively. An average of 5 colonies were imaged in each condition for each mutant.

The fifth experimental batch is presented in Section “Impact of pre-2×SLAD and on invasive growth in gene-deletion mutants of AWRI 796”. It consisted of the AWRI 796 strain and yeast gene deletants: *alr2, dur3, nrt1, tmn3, pma2, put4, sac3*. The experimental design for each strain/mutant was a (2 × 2 × 2 = 8) fully factorial design varying ammonium sulfate (50 or 75 µM), sodium sulfide (0 or 400 µM) and plates washed on day (4 or 6). All conditions used pre-2×SLAD with a 2×SLAD BD agar. On average, there were 39 colonies per plate before washing and 30 colonies that were growing invasively. An average of 6 colonies were imaged in each condition for each mutant.

### Statistical Analysis

All data analyses were conducted in the statistical computing environment R^33^ using the RStudio graphical user interface^34^. Data manipulation and visualisation were performed primarily with the tidyverse^35^, including the ggplot2 package for graphics^36^. Two outcome types were considered in this study. The first was the number of colonies on a given plate that exhibited any invasive growth (presence of invasion). The second was the extent of invasive growth remaining after washing (degree of invasion).

A natural statistical approach for analysing the presence of invasion data was logistic regression, modelling the number of invasive colonies relative to the total number of colonies on each plate. However, the proportion of colonies showing invasive growth after washing was frequently either zero or one (e.g., Figure 4). This complete or near-complete separation resulted in convergence difficulties for logistic regression models. An alternative approach would be Poisson regression with an exposure term for the total number of colonies per plate, but this method encountered similar limitations due to the extreme distribution of outcomes. Consequently, we relied on exploratory data analysis to summarise and visualise patterns in the presence of invasion, rather than formal statistical testing for this outcome.

We used a Beta regression model to analyse the degree of invasion values *D*_*i, j*_, where *D*_*i, j*_ denotes the measured degree of invasion for colony *i* under experimental condition *j*. This approach is appropriate for continuous outcomes bounded within the (0, 1) interval and assumed to follow a Beta(*µ, φ*) distribution, where *µ* denotes the mean of the outcome and *φ* the precision parameter. For a fixed *µ*, larger values of *φ* correspond to lower variance. The model was implemented in R using the `betareg` package^37^, assuming constant precision across observations. The mean parameter *µ* was modelled using a logit link as a function of experimental predictors and their interactions where relevant, as determined by the study design. As the Beta distribution is defined only on the open interval (0, 1), colonies with zero invasive growth were adjusted by adding a small constant (0.001) prior to modelling. This adjustment reflected the practical detection limit of the image analysis procedure, which cannot resolve arbitrarily small amounts of invasion. A summary of the regression models is provided in Table 2. Coefficient estimates for the *µ* parameter are presented in Figures 5-11 but full model parameter estimate tables are also included in the Supplementary materials.

**Table 1.**
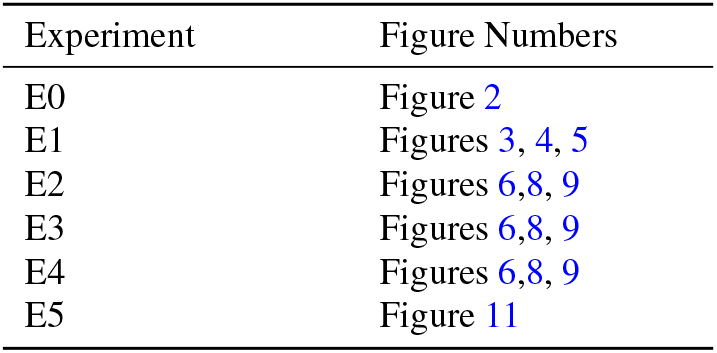
Summary of experimental batch (E0-E5) and their association to each experiment with the figure produced in the main manuscript.

**Table 2.**
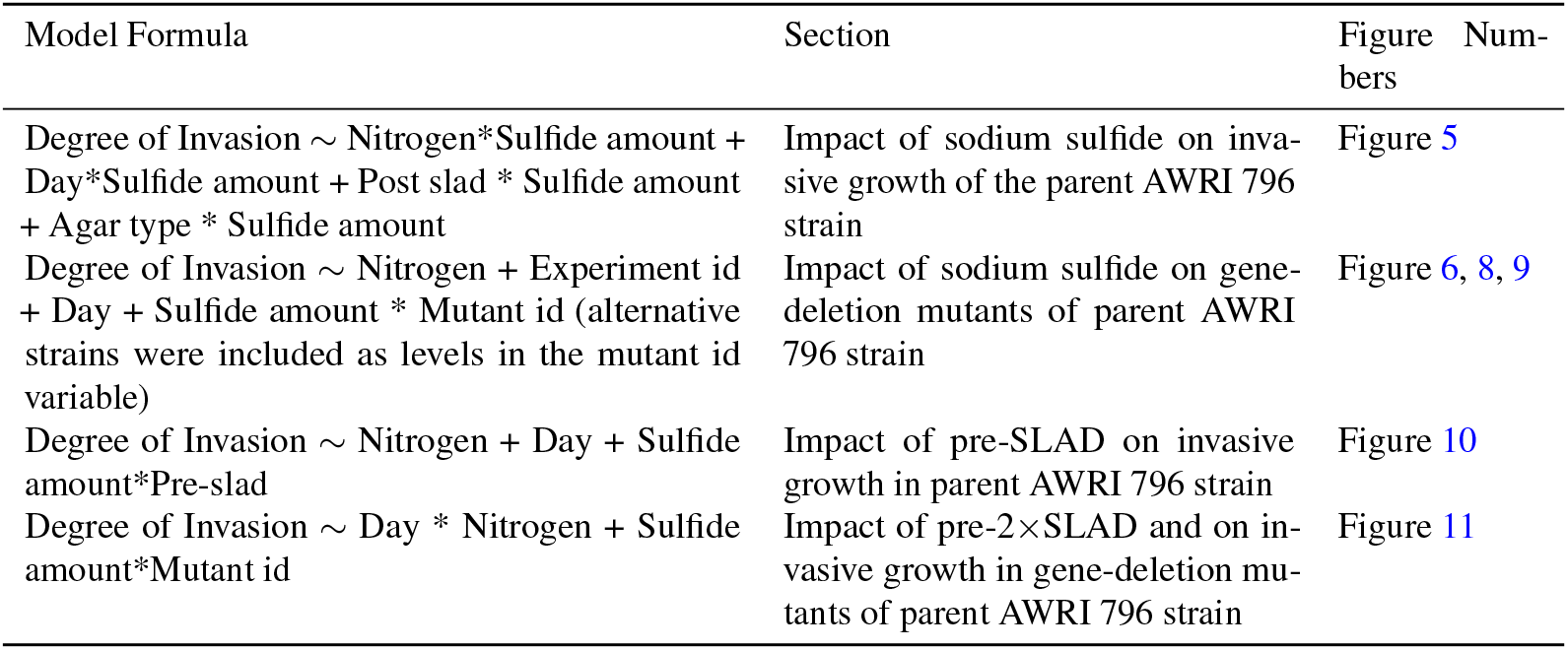
Summary of regression models and the relevant sections, figures and supplementary materials describing the estimates. Regression models are specified using R formula shorthand, where a var1*var2 indicates main effects for var 1, var 2 and the interaction between var 1 and var 2 will be included. Section refers to the main (where results and coefficient estimates for *µ* parameter are presented) and the supplementary materials (where full model coefficient estimates are included in tables).

For all experiments except the first (which examines nutrient effects alone), our primary parameter of interest was the slope associated with sodium sulfide, representing the change in invasive growth between the absence and presence of sodium sulfide. Because zero sodium sulfide was specified as the reference level, this coefficient could be interpreted directly as the effect of adding sodium sulfide. In the analyses of gene-deletion mutants and additional yeast strains, the parent strain AWRI 796 was used as the reference class, allowing the parameters for other strains to be interpreted relative to the parent. All remaining categorical variables were coded using their default reference levels: day 3 for plate washing time, 1×SLAD for medium strength, and 50 µM ammonium sulfate for nitrogen concentration.

## Acknowledgements

We acknowledge funding from the Australian Research Council (grant numbers DP230100406, DE240100097, and IC170100008) and Wine Australia (UA1803_2.1).

## Additional information

GitHub link to data and code: https://github.com/kaili2019/Li2026-Invasive-Growth. Original TIFF image data is available upon reasonable request from the corresponding author.

