## Supplementary Material for "Experimental design and analysis of sulfide-induced invasive growth in wine yeast"

### ABSTRACT

Yeast's ability to invade surfaces has important implications for infections and food contamination. Invasive growth in yeast is influenced by genetic and environmental factors. In this exploratory study, we investigated the effects of sodium sulfide, gene deletions, and environmental conditions on the invasive behaviour of the wine yeast strain AWRI 796. Sodium sulfide enhanced invasion in the (parent) AWRI 796 strain under nitrogen-limiting conditions, although its effect was obscured by experimental variability and pre-culture conditions. Genetic factors had a major effect on the overall invasive phenotype, with deletion of key genes suppressing invasion. Most gene-deletion mutants did not significantly affect how the colony responded to sulfide. In addition to sulfide and genotype, environmental conditions also influenced invasive behaviour. The pre-2$\times$SLAD pre-culture condition was best for detecting sulfide-induced growth, and later plate washing time and decreased nutrient levels enhanced invasiveness. Our experimental design and findings provide a framework for understanding the determinants of yeast invasiveness, which may inform future studies on filamentous yeast behaviour.

### Supplementary Material

#### Data pre-processing and cleaning

The first task was to pre-process the image data in MATLAB into binary images to compute the surface area and invasive area after washing. The image dataset was provided in one of two formats. The first format was uint8 images, which we directly processed using TAMMiCol. The second format was uint16 images, which required contrast adjustment and conversion to uint8 before these images were processed in TAMMiCol. After TAMMiCol had binarised all the images, we computed the area of the surface and invasive area after washing. From this, we also computed the ratio of the invasive area to surface area as a relative measure of invasion across the surface. Each image had a unique name detailing its growth conditions and strain in the format

yeast/mutant ID _ Nutrient type and concentration _ liquid starter culture medium _ experimental medium _ experimental time (days) _ ID for replicates in each condition.

We extracted all the information from the file names of each image, and together with their respective area data, we tabulated the data into CSV files. This was done once for each experiment (initially denoted A-E before being redefined into numeric E1-E5 representing the first to last completed), resulting in five CSV files, which were later further processed in R.

The count data were provided in Excel spreadsheets. We first read the sheets into R and saved them as CSV files. Then, we cleaned each CSV file by extracting the rows that contained the count information, removing completely empty rows and converting the file name (which described experimental conditions/manipulations) into additional variables. Dates were used to confirm the recorded number of days between when colonies were inoculated and when they were washed. Finally, specific columns were selected from each individual CSV file containing data from each sheet. These were yeast strain, liquid starter culture medium, experimental medium, nutrient level and concentration, amount of sodium sulfide, total colonies count before wash, total invasive colonies after wash, agar type, experiment id (E1 through E5), sheet day and parent or not.

After the image and count data were combined into their individual CSVs, we cleaned to ensure they had consistent naming across the two datasets. All the file names from the image dataset and count dataset were first renamed to have the same variable names for entities that shared the same attribute. These were agar type, nutrient, days of experiment, liquid starter culture medium, experimental medium, amount of sodium sulfide, experiment order and mutant ID.

When the colonies were imaged, there were some images that were not taken. For transparency, we report the total number of missing unwashed and missing washed images per experiment. Missing either image meant that we could not calculate the degree of invasion and so that colony was excluded from our analysis. We tabulated all of the missing images in Tables 1,2,3,4,5,6. The 'Experiment' column is the order of the experiment. The 'Identifier' column is either the parent strain AWRI 796, which we denote 'Parent', or otherwise is a mutant deletion gene with a given abbreviation.

**Table 1.** Summary of missing images from experiment E0.

| Experiment | Identifier | Sulfide Amount (µM) | Day | Nutrient Level (µM) | pre SLAD | post SLAD | # Missing Washed Images | # Missing Unwashed Images |
|---|---|---|---|---|---|---|---|---|
| E0 | AWRI 796 | 0 | 10 | BD500 | 1× | 2× | 1 | 1 |

**Table 2.** Summary of missing images from experiment E1.

| Experiment | Identifier | Sulfide Amount (µM) | Day | Nutrient Level (µM) | pre SLAD | post SLAD | # Missing Washed Images | # Missing Unwashed Images |
|---|---|---|---|---|---|---|---|---|
| E1 | AWRI 796 | 750 | 3 | BD50 | 1× | 2× | 0 | 2 |
| E1 | AWRI 796 | 0 | 6 | BD50 | 1× | 2× | 0 | 1 |
| E1 | AWRI 796 | 750 | 6 | BD75 | 1× | 2× | 1 | 0 |
| E1 | AWRI 796 | 750 | 6 | BD75 | 1× | 2× | 0 | 5 |
| E1 | AWRI 796 | 750 | 6 | BD100 | 1× | 2× | 0 | 5 |
| E1 | AWRI 796 | 0 | 6 | BD50 | 1× | 2× | 0 | 1 |
| E1 | AWRI 796 | 400 | 6 | Ox50 | 1× | 2× | 0 | 5 |
| E1 | AWRI 796 | 750 | 6 | Ox50 | 1× | 2× | 0 | 5 |
| E1 | AWRI 796 | 0 | 6 | Ox75 | 1× | 2× | 0 | 5 |
| E1 | AWRI 796 | 750 | 6 | Ox75 | 1× | 2× | 0 | 5 |
| E1 | AWRI 796 | 100 | 6 | Ox100 | 1× | 2× | 0 | 5 |
| E1 | AWRI 796 | 400 | 6 | Ox100 | 1× | 2× | 0 | 6 |
| E1 | AWRI 796 | 750 | 6 | Ox100 | 1× | 2× | 0 | 5 |

**Table 3.** Summary of missing images from experiment E2.

| Experiment | Identifier | Sulfide Amount (µM) | Day | Nutrient Level (µM) | pre SLAD | post SLAD | # Missing Washed Images | # Missing Unwashed Images |
|---|---|---|---|---|---|---|---|---|
| E2 | AWRI 796 | 400 | 3 | BD50 | 1× | 2× | 4 | 0 |
| E2 | AWRI 796 | 0 | 3 | BD50 | 1× | 2× | 2 | 0 |
| E2 | AWRI 796 | 400 | 3 | BD50 | 1× | 2× | 0 | 1 |
| E2 | *fat1* | 0 | 3 | BD75 | 1× | 2× | 1 | 0 |
| E2 | *y0r1* | 400 | 3 | BD50 | 1× | 2× | 1 | 0 |
| E2 | L2056 | 400 | 3 | BD50 | 1× | 2× | 2 | 0 |
| E2 | L2056 | 400 | 3 | BD75 | 1× | 2× | 4 | 0 |
| E2 | *skp2* | 400 | 3 | BD50 | 1× | 2× | 0 | 4 |
| E2 | *skp2* | 0 | 3 | BD50 | 1× | 2× | 0 | 4 |
| E2 | *skp2* | 400 | 3 | BD75 | 1× | 2× | 0 | 5 |
| E2 | *skp2* | 0 | 3 | BD75 | 1× | 2× | 0 | 4 |
| E2 | *soa1* | 0 | 3 | BD50 | 1× | 2× | 0 | 1 |
| E2 | *gup1* | 400 | 3 | BD50 | 1× | 2× | 0 | 3 |
| E2 | *gup1* | 400 | 3 | BD75 | 1× | 2× | 0 | 4 |
| E2 | *gup1* | 0 | 3 | BD75 | 1× | 2× | 0 | 4 |
| E2 | AWRI 796 | 400 | 6 | BD50 | 1× | 2× | 4 | 1 |
| E2 | AWRI 796 | 0 | 6 | BD50 | 1× | 2× | 4 | 1 |
| E2 | AWRI 796 | 400 | 6 | BD75 | 1× | 2× | 4 | 1 |
| E2 | AWRI 796 | 0 | 6 | BD75 | 1× | 2× | 4 | 1 |
| E2 | *gup1* | 400 | 6 | BD50 | 1× | 2× | 0 | 1 |
| E2 | *skp2* | 0 | 6 | BD50 | 1× | 2× | 0 | 1 |
| E2 | Σ1278b | 0 | 6 | BD50 | 1× | 2× | 0 | 1 |
| E2 | Σ1278b | 400 | 6 | BD75 | 1× | 2× | 0 | 1 |
| E2 | *fat1* | 0 | 6 | BD75 | 1× | 2× | 0 | 2 |
| E2 | *fat1* | 400 | 6 | BD50 | 1× | 2× | 0 | 1 |
| E2 | *nrt1* | 0 | 6 | BD75 | 1× | 2× | 0 | 4 |
| E2 | *soa1* | 400 | 6 | BD50 | 1× | 2× | 0 | 1 |
| E2 | L2056 | 0 | 6 | BD75 | 1× | 2× | 0 | 6 |
| E2 | L2056 | 0 | 6 | BD50 | 1× | 2× | 0 | 1 |
| E2 | L2056 | 400 | 6 | BD75 | 1× | 2× | 0 | 4 |

**Table 4.** Summary of missing images from experiment E3.

| Experiment | Identifier | Sulfide Amount (µM) | Day | Nutrient Level (µM) | pre SLAD | post SLAD | # Missing Washed Images | # Missing Unwashed Images |
|---|---|---|---|---|---|---|---|---|
| E3 | *ccz1* | 0 | 4 | BD50 | 1× | 2× | 0 | 1 |
| E3 | *ccz1* | 400 | 4 | BD50 | 1× | 2× | 0 | 1 |
| E3 | *cdh1* | 400 | 4 | BD50 | 1× | 2× | 0 | 1 |
| E3 | *rps8* | 0 | 4 | BD50 | 1× | 2× | 1 | 0 |
| E3 | *whi3* | 400 | 4 | BD50 | 1× | 2× | 4 | 0 |
| E3 | *whi3* | 0 | 4 | BD50 | 1× | 2× | 4 | 0 |
| E3 | *whi3* | 400 | 4 | BD75 | 1× | 2× | 4 | 0 |
| E3 | *whi3* | 0 | 4 | BD75 | 1× | 2× | 4 | 0 |
| E3 | *ccz1* | 0 | 6 | BD50 | 1× | 2× | 1 | 0 |
| E3 | *cdh1* | 400 | 6 | BD50 | 1× | 2× | 2 | 0 |
| E3 | *cdh1* | 0 | 6 | BD50 | 1× | 2× | 1 | 0 |
| E3 | *cdh1* | 400 | 6 | BD75 | 1× | 2× | 2 | 0 |
| E3 | *vps28* | 400 | 6 | BD50 | 1× | 2× | 1 | 0 |
| E3 | *vps28* | 0 | 6 | BD50 | 1× | 2× | 1 | 0 |
| E3 | *whi3* | 400 | 6 | BD50 | 1× | 2× | 0 | 4 |
| E3 | *whi3* | 0 | 6 | BD50 | 1× | 2× | 0 | 4 |
| E3 | *whi3* | 400 | 6 | BD75 | 1× | 2× | 0 | 4 |
| E3 | *whi3* | 0 | 6 | BD75 | 1× | 2× | 0 | 4 |

**Table 5.** Summary of missing images from experiment E4.

| Experiment | Identifier | Sulfide Amount (µM) | Day | Nutrient Level (µM) | pre SLAD | post SLAD | # Missing Washed Images | # Missing Unwashed Images |
|---|---|---|---|---|---|---|---|---|
| E4 | *ato3* | 400 | 4 | BD75 | 1× | 2× | 0 | 1 |
| E4 | *ato3* | 0 | 4 | BD75 | 1× | 2× | 0 | 1 |
| E4 | *top4* | 400 | 4 | BD50 | 1× | 2× | 0 | 3 |
| E4 | AWRI 796 | 400 | 6 | BD75 | 2× | 2× | 0 | 2 |
| E4 | *alr2* | 400 | 6 | BD50 | 1× | 2× | 0 | 4 |
| E4 | *alr2* | 0 | 6 | BD50 | 1× | 2× | 0 | 1 |
| E4 | *alr2* | 0 | 6 | BD75 | 1× | 2× | 0 | 1 |
| E4 | *ato3* | 0 | 6 | BD50 | 1× | 2× | 0 | 1 |
| E4 | *msb2* | 400 | 6 | BD50 | 1× | 2× | 0 | 1 |

**Table 6.** Summary of missing images from experiment E5.

| Experiment | Identifier | Sulfide Amount (µM) | Day | Nutrient Level (µM) | pre SLAD | post SLAD | # Missing Washed Images | # Missing Unwashed Images |
|---|---|---|---|---|---|---|---|---|
| E5 | *put4* | 400 | 4 | BD50 | 2× | 2× | 4 | 0 |
| E5 | *put4* | 0 | 4 | BD50 | 2× | 2× | 5 | 0 |
| E5 | *put4* | 400 | 4 | BD75 | 2× | 2× | 3 | 0 |
| E5 | *put4* | 0 | 4 | BD75 | 2× | × | 2 | 0 |
| E5 | *yhl008c* | 400 | 4 | BD50 | 2× | 2× | 1 | 0 |
| E5 | *yhl008c* | 0 | 4 | BD50 | 2× | 2× | 1 | 0 |
| E5 | *yhl008c* | 0 | 4 | BD75 | 2× | 2× | 2 | 0 |
| E5 | *yhl008c* | 400 | 4 | BD50 | 2× | 2× | 1 | 0 |
| E5 | *alr2* | 400 | 4 | BD75 | 2× | 2× | 1 | 0 |
| E5 | *nrt1* | 0 | 4 | BD50 | 2× | 2× | 1 | 0 |
| E5 | *sac3* | 400 | 4 | BD50 | 2× | 2× | 0 | 4 |
| E5 | *sac3* | 0 | 4 | BD50 | 2× | 2× | 0 | 4 |
| E5 | *sac3* | 400 | 4 | BD75 | 2× | 2× | 0 | 4 |
| E5 | *sac3* | 0 | 4 | BD75 | 2× | 2× | 0 | 4 |
| E5 | *vht1* | 0 | 4 | BD75 | 2× | 2× | 0 | 2 |
| E5 | *put4* | 400 | 4 | BD50 | 2× | 2× | 0 | 2 |
| E5 | *put4* | 0 | 4 | BD50 | 2× | 2× | 0 | 1 |
| E5 | *put4* | 0 | 6 | BD50 | 2× | 2× | 3 | 0 |
| E5 | *nrt1* | 400 | 6 | BD50 | 2× | 2× | 1 | 0 |
| E5 | *sac3* | 400 | 6 | BD50 | 2× | 2× | 0 | 6 |
| E5 | *sac3* | 400 | 6 | BD75 | 2× | 2× | 0 | 3 |
| E5 | *sac3* | 0 | 6 | BD75 | 2× | 2× | 0 | 2 |
| E5 | *vht1* | 400 | 6 | BD75 | 2× | 2× | 0 | 4 |
| E5 | *put4* | 0 | 6 | BD50 | 2× | 2× | 0 | 1 |

### Statistical results

#### *Impact of sodium sulfide on invasive growth of the parent AWRI 796 strain*

A regression analysis predicting degree of invasion with main effects for the ammonium sulfate concentration, sodium sulfide level, day of washing, the SLAD concentration and manufacturer plus interactions for each with the sodium sulfide level. Raw data for both presence and degree of invasion are presented in the main text. Full regression model estimates are included in Table 7.

**Table 7.** Full parameter estimates of model predicting the ratio of washed surface colony area to unwashed surface colony area in Experiment 1. Parameter estimates for $\mu$ shown in Figure 5 of the main text. This Beta regression was fit with main effects for all relevant experimental conditions with interactions between experimental conditions and the sodium sulfide intervention. Zero sodium sulfide was the sodium sulfide reference category, with other reference categories as BD Agar, 1×SLAD, Day 3 and 50 µM of ammonium sulfate.

| | Est. | S.E. | t | p |
|---|---|---|---|---|
| sodium sulfide 400 µM: Oxoid agar | −0.933 | 0.260 | −3.585 | <0.001 |
| sodium sulfide 400 µM: 2×SLAD | 0.231 | 0.253 | 0.915 | 0.360 |
| sodium sulfide 400 µM: day6 | −0.248 | 0.249 | −0.995 | 0.320 |
| sodium sulfide 400 µM: ammonium sulfate 75 µM | −1.093 | 0.284 | −3.855 | <0.001 |
| sodium sulfide 400 µM: ammonium sulfate 100 µM | −0.943 | 0.281 | −3.355 | <0.001 |
| sodium sulfide 750 µM: Oxoid agar | −0.692 | 0.274 | −2.525 | 0.012 |
| sodium sulfide 750 µM: 2×SLAD | −0.217 | 0.276 | −0.785 | 0.432 |
| sodium sulfide 750 µM: day6 | 0.451 | 0.266 | 1.698 | 0.089 |
| sodium sulfide 750 µM: ammonium sulfate 75 µM | −2.189 | 0.290 | −7.552 | <0.001 |
| sodium sulfide 750 µM: ammonium sulfate 100 µM | −1.975 | 0.286 | −6.895 | <0.001 |
| Oxoid agar | 2.583 | 0.199 | 12.968 | <0.001 |
| 2×SLAD | 2.210 | 0.185 | 11.960 | <0.001 |
| Day 6 | 2.143 | 0.187 | 11.485 | <0.001 |
| 75 µM ammonium sulfate | 2.143 | 0.207 | 10.349 | <0.001 |
| 100 µM ammonium sulfate | 1.387 | 0.209 | 6.654 | <0.001 |
| 400 µM sodium sulfide | 1.548 | 0.347 | 4.459 | <0.001 |
| 750 µM sodium sulfide | 2.684 | 0.361 | 7.444 | <0.001 |
| Intercept | −7.199 | 0.278 | −25.906 | <0.001 |
| $\phi$ | 8.097 | 0.709 | 11.414 | <0.001 |
| Num.Obs. | 336 | | | |
| R2 | 0.695 | | | |
| R2 Pseudo | 0.695 | | | |
| AIC | −1322.3 | | | |
| BIC | −1249.7 | | | |
| Log.Lik. | 680.132 | | | |
| RMSE | 0.10 | | | |
| DF Null | 334 | | | |
| DF Resid | 317 | | | |

#### *Impact of sodium sulfide on gene-deletion mutants of parent AWRI 796 strain*

A regression analysis predicting degree of invasion with main effects for the ammonium sulfate concentration, experimental id, sodium sulfide level, day of washing and the mutant or strain. An interaction was also included for each mutant or strain with the sodium sulfide level (AWRI 796 included as the reference class). Raw data of the degree of invasion for the mutations from AWRI 796 are presented in Figure 1 of supplementary materials. Count data is included in the Github repository. Full regression model estimates are included in Table 8.

**Table 8.** Full parameter estimates of model predicting the ratio of washed surface colony area to unwashed surface colony area for mutants and alternative strains in experiments 2, 3 and 4. Parameter estimates for $\mu$ shown in Figures 6, 8 and 9 in the main text. This Beta regression was fit with main effects for all relevant experimental conditions, the strain/mutant and interactions between the strain/mutant and the sodium sulfide intervention. Zero sodium sulfide was the sodium sulfide reference category, with other reference categories as Day 3, 50 µM of ammonium sulfate, first experimental batch and parent yeast strain 796.

| | Est. | S.E. | t | p |
|---|---|---|---|---|
| Sodium Sulfide 400 $\mu$M: yor1 | 0.581 | 0.313 | 1.856 | 0.063 |
| Sodium Sulfide 400 $\mu$M: alr2 | −0.134 | 0.341 | −0.393 | 0.695 |
| Sodium Sulfide 400 $\mu$M: ato3 | 0.144 | 0.340 | 0.424 | 0.672 |
| Sodium Sulfide 400 $\mu$M: ccz1 | 0.185 | 0.292 | 0.635 | 0.526 |
| Sodium Sulfide 400 $\mu$M: cdh1 | −0.311 | 0.341 | −0.912 | 0.362 |
| Sodium Sulfide 400 $\mu$M: cvt16 | 0.454 | 0.288 | 1.577 | 0.115 |
| Sodium Sulfide 400 $\mu$M: dur3 | −0.425 | 0.319 | −1.334 | 0.182 |
| Sodium Sulfide 400 $\mu$M: fat1 | 0.316 | 0.326 | 0.967 | 0.334 |
| Sodium Sulfide 400 $\mu$M: fui1 | 0.465 | 0.335 | 1.387 | 0.166 |
| Sodium Sulfide 400 $\mu$M: gup1 | −0.118 | 0.394 | −0.299 | 0.765 |
| Sodium Sulfide 400 $\mu$M: mid1 | 0.040 | 0.319 | 0.126 | 0.900 |
| Sodium Sulfide 400 $\mu$M: msa1 | 0.080 | 0.288 | 0.278 | 0.781 |
| Sodium Sulfide 400 $\mu$M: msb2 | −0.278 | 0.341 | −0.814 | 0.416 |
| Sodium Sulfide 400 $\mu$M: nrt1 | 1.552 | 0.320 | 4.858 | <0.001 |
| Sodium Sulfide 400 $\mu$M: pep12 | 0.355 | 0.291 | 1.220 | 0.223 |
| Sodium Sulfide 400 $\mu$M: rps8a | 0.443 | 0.334 | 1.328 | 0.184 |
| Sodium Sulfide 400 $\mu$M: skp2 | −0.315 | 0.429 | −0.734 | 0.463 |
| Sodium Sulfide 400 $\mu$M: soa1 | 0.294 | 0.315 | 0.933 | 0.351 |
| Sodium Sulfide 400 $\mu$M: tmn3 | −0.093 | 0.296 | −0.313 | 0.754 |
| Sodium Sulfide 400 $\mu$M: tpo4 | −0.396 | 0.288 | −1.374 | 0.169 |
| Sodium Sulfide 400 $\mu$M: vps28 | −0.315 | 0.341 | −0.923 | 0.356 |
| Sodium Sulfide 400 $\mu$M: whi3 | −0.386 | 0.688 | −0.561 | 0.575 |

| | | | | |
|---|---|---|---|---|
| Sodium Sulfide 400 $\mu$M: Σ1278b | −0.234 | 0.299 | −0.784 | 0.433 |
| Sodium Sulfide 400 $\mu$M: L2056 | 1.704 | 0.298 | 5.714 | <0.001 |
| yor1 | −0.368 | 0.262 | −1.405 | 0.160 |
| alr2 | −1.324 | 0.256 | −5.165 | <0.001 |
| ato3 | −1.254 | 0.256 | −4.896 | <0.001 |
| ccz1 | −0.670 | 0.226 | −2.972 | 0.003 |
| cdh1 | −2.698 | 0.253 | −10.646 | <0.001 |
| cvt16 | −0.217 | 0.218 | −0.995 | 0.320 |
| dur3 | 0.245 | 0.239 | 1.021 | 0.307 |
| fat1 | −0.970 | 0.264 | −3.677 | <0.001 |
| fui1 | −1.060 | 0.255 | −4.152 | <0.001 |
| gup1 | −1.918 | 0.297 | −6.463 | <0.001 |
| mid1 | 0.005 | 0.244 | 0.020 | 0.984 |
| msa1 | −0.180 | 0.217 | −0.828 | 0.408 |
| msb2 | −1.324 | 0.256 | −5.165 | <0.001 |
| nrt1 | −1.779 | 0.266 | −6.701 | <0.001 |
| pep12 | −0.429 | 0.221 | −1.938 | 0.053 |
| rps8a | −2.327 | 0.252 | −9.246 | <0.001 |
| skp2 | −2.156 | 0.332 | −6.489 | <0.001 |
| soa1 | −0.200 | 0.264 | −0.757 | 0.449 |
| tmn3 | −0.527 | 0.223 | −2.366 | 0.018 |
| tpo4 | 1.431 | 0.221 | 6.480 | <0.001 |
| vps28 | −2.726 | 0.254 | −10.752 | <0.001 |
| whi3 | −2.655 | 0.466 | −5.695 | <0.001 |
| Σ1278b | 1.995 | 0.248 | 8.055 | <0.001 |
| L2056 | −1.665 | 0.258 | −6.461 | <0.001 |
| 75 $\mu$M Ammonium Sulfate | 0.079 | 0.055 | 1.442 | 0.149 |
| Day 6 | 1.859 | 0.095 | 19.601 | <0.001 |
| Day 4 | 1.213 | 0.112 | 10.809 | <0.001 |
| Third batch | −0.713 | 0.173 | −4.115 | <0.001 |
| Second batch | 0.690 | 0.177 | 3.890 | <0.001 |
| 400 $\mu$M Sodium Sulfide | 0.315 | 0.120 | 2.619 | 0.009 |
| Intercept | −2.800 | 0.170 | −16.452 | <0.001 |

| | | | | |
|---|---|---|---|---|
| $\phi$ | 5.023 | 0.235 | 21.368 | <0.001 |
| Num.Obs. | 1132 | | | |
| R2 | 0.693 | | | |
| R2 Pseudo | 0.693 | | | |
| AIC | −4196.2 | | | |
| BIC | −3914.5 | | | |
| Log.Lik. | 2154.120 | | | |
| RMSE | 0.13 | | | |
| DF Null | 1130 | | | |
| DF Resid | 1076 | | | |

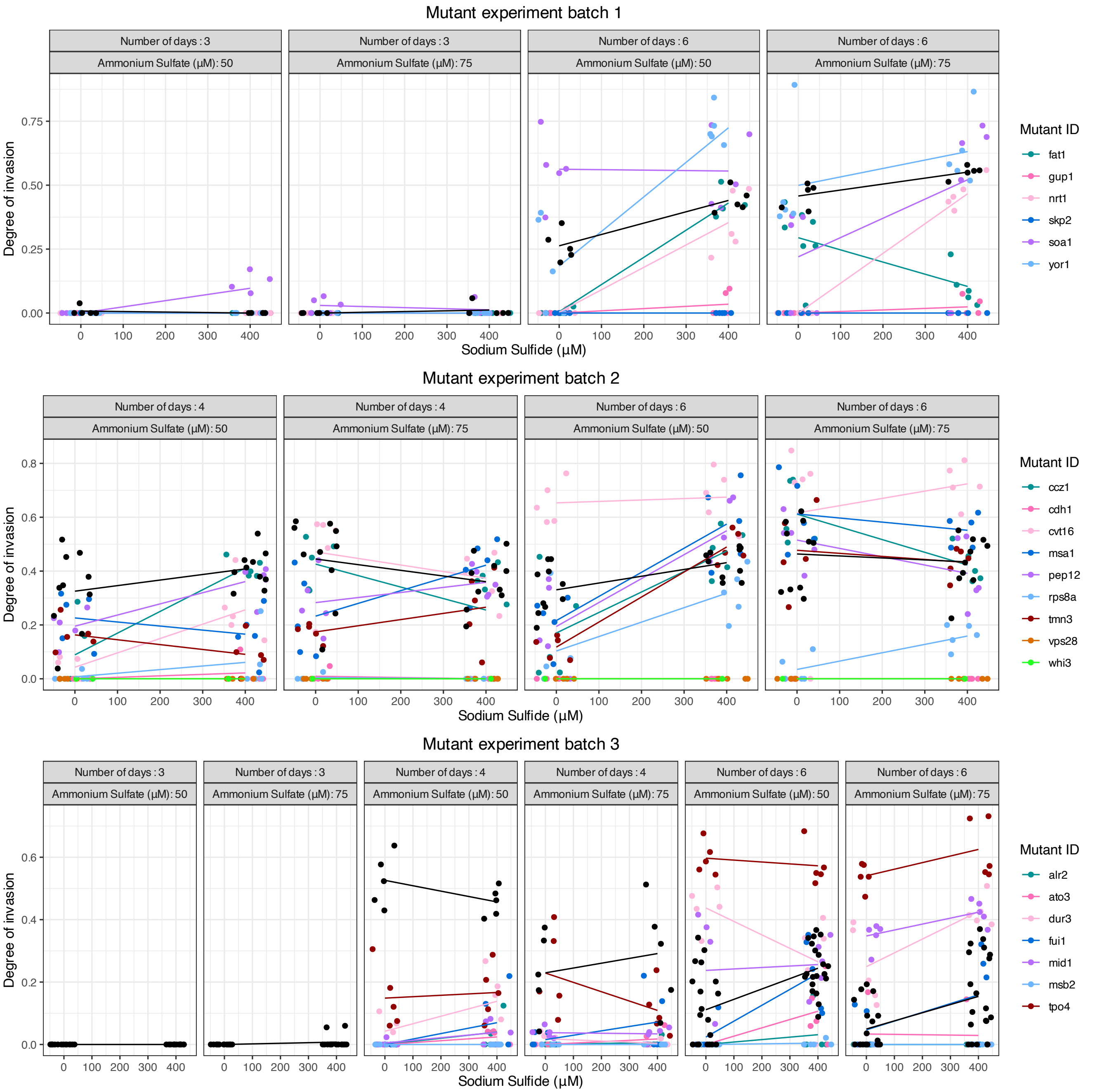


**Figure 1.** Raw ratio of washed to unwashed surface area for the mutant experiments on a 2×SLAD. Black is used for the parent yeast, with points representing individual colonies. Points are jittered on the *x*-axis to reduce overplotting - sodium sulfide levels are only 0 and 400. A line joining the average for a particular mutant with zero sodium sulfide and 400 μM sodium sulfide has been drawn to help to visually identify mutants response to sodium sulfide. Note the colour mapping to mutant is reused on each row (corresponding to experiment batch) to allow the colours to be sufficiently accessible and distinguishable. As we can see, there is quite a lot of variability within a mutant in a experimental condition, suggesting more observations per category are needed. Mutants with slopes that are negative or less steep than the corresponding parent (black) slope might be of future experimental interest.

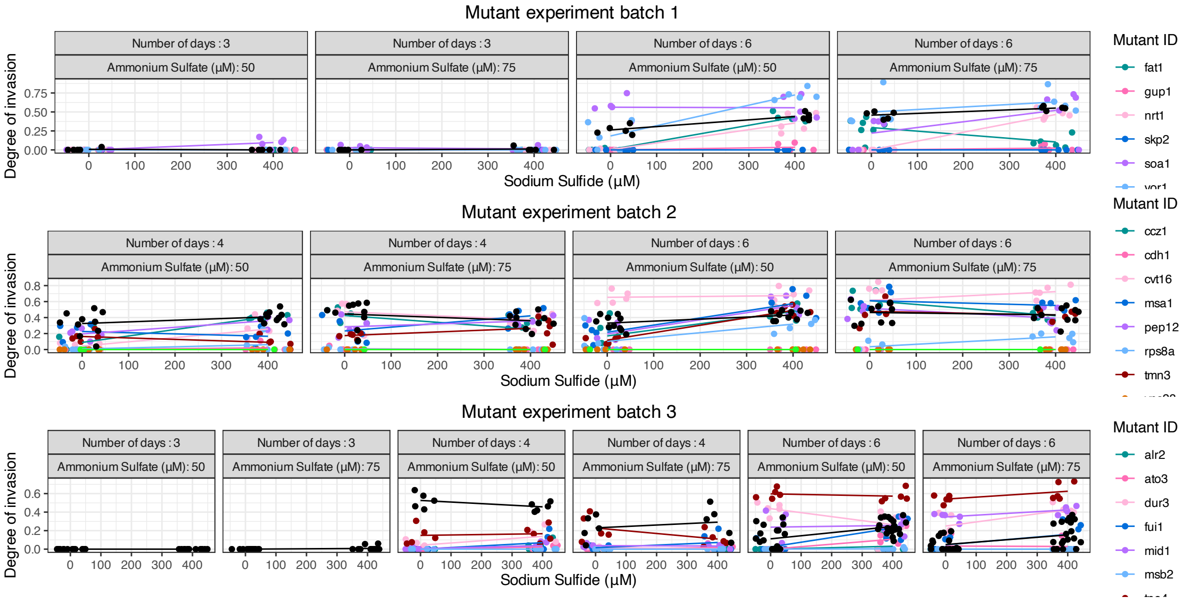


**Figure 2.** Measurement of yeast invasion using image analysis (top row) and count data (bottom row). The x-axis represents the amount of sodium sulfide present, and hence the line in the top row represents the difference between no-sulfur and sulfur conditions for the strain AWRI 796 (orange, light grey), strain Σ 1278b (pink, medium grey) and strain L2056 (purple, dark grey). The line is draw between the average values of yeast image for that particular experimental condition. The bottom row represents the proportion of yeast colonies that grew invasively across the whole plate, as counted in the lab.

#### ***Impact of pre-SLAD on invasive growth in parent AWRI 796 strain***

A regression analysis predicting degree of invasion with main effects for the ammonium sulfate concentration, sodium sulfide level, day of washing and the pre-SLAD treatment. An interaction was also included for the pre-SLAD treatment. Raw data of the degree of invasion for the mutations from AWRI 796 are presented in Figure 3 of supplementary materials. Full regression model estimates are included in Table 9.

**Table 9.** Full parameter estimates of model predicting the ratio of washed surface colony area to unwashed surface colony area for parents with different SLADs in experiment 4. Parameter estimates for $\mu$ shown in Figure 10 of the main text. This Beta regression was fit with main effects for all relevant experimental conditions and an interaction between the pre-SLAD and the sodium sulfide intervention. Zero sodium sulfide was the sodium sulfide reference category, with other reference categories as Day 3, 50 µM of ammonium sulfate and 1 1×SLAD.

| | Est. | S.E. | t | p |
|---|---|---|---|---|
| sodium sulfide 750 µM: pre-2×SLAD | 0.667 | 0.258 | 2.588 | 0.010 |
| pre-2×SLAD | −0.419 | 0.203 | −2.069 | 0.039 |
| Day 6 | 1.336 | 0.154 | 8.652 | <0.001 |
| Day 4 | 2.739 | 0.195 | 14.080 | <0.001 |
| 75 µM ammonium sulfate | −0.622 | 0.122 | −5.085 | <0.001 |
| 400 µM sodium sulfide | 0.348 | 0.150 | 2.327 | 0.020 |
| Intercept | −3.171 | 0.177 | −17.928 | <0.001 |
| $\phi$ | 12.306 | 1.508 | 8.158 | <0.001 |
| Num.Obs. | 160 | | | |
| R2 | 0.637 | | | |
| R2 Pseudo | 0.637 | | | |
| AIC | −511.2 | | | |
| BIC | −486.6 | | | |
| Log.Lik. | 263.579 | | | |
| RMSE | 0.09 | | | |
| DF Null | 158 | | | |
| DF Resid | 152 | | | |

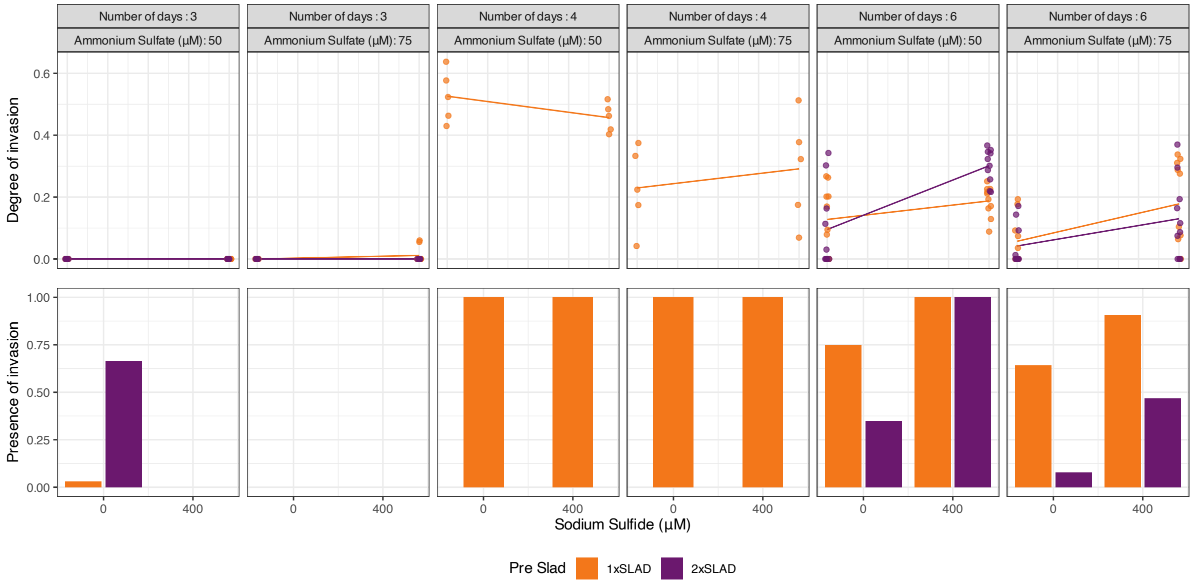


**Figure 3.** Measurement of yeast invasion using image analysis (top row) and count data (bottom row). The x-axis represents the amount of sodium sulfide present, and hence the line in the top row represents the difference between no-sulfur and sulfur conditions for the pre-1×SLAD (orange, light grey) and pre-2×SLAD (purple, dark grey). The line is draw between the average values of yeast image for that particular experimental condition. The bottom row represents the proportion of yeast colonies that grew invasively across the whole plate, as counted in the lab.

#### *Impact of pre-2×SLAD and on invasive growth in gene-deletion mutants of parent AWRI 796 strain*

A regression analysis predicting degree of invasion with main effects for the ammonium sulfate concentration, experimental id, sodium sulfide level, day of washing and the mutant or strain using the pre-1×SLAD. An interaction was also included for each mutant or strain with the sodium sulfide level (AWRI 796 included as the reference class). Raw data of the degree of invasion for the mutations from AWRI 796 are presented in Figure 4 of supplementary materials. Full regression model estimates are included in Table 10.

**Table 10.** Full parameter estimates of model predicting the ratio of washed surface colony area to unwashed surface colony area for mutants and alternative strains in experiments 5 with the 2×SLAD liquid start culture medium. Parameter estimates for $\mu$ shown in Figure 11 of the main manuscript. This Beta regression was fit with main effects for all relevant experimental conditions, the mutant and interactions between the mutant and the sodium sulfide intervention. Zero sodium sulfide was the sodium sulfide reference category, with other reference categories as Day 3, 50 µM of ammonium sulfate, first experimental batch and parent yeast strain 796.

| | Est. | S.E. | t | p |
|---|---|---|---|---|
| Sodium Sulfide 400 $\mu$M: yhl008c | 0.715 | 0.254 | 2.815 | 0.005 |
| Sodium Sulfide 400 $\mu$M: u0394tmn3 | 0.448 | 0.291 | 1.538 | 0.124 |
| Sodium Sulfide 400 $\mu$M: put4 | 1.416 | 0.306 | 4.634 | <0.001 |
| Sodium Sulfide 400 $\mu$M: pma2 | 0.486 | 0.292 | 1.664 | 0.096 |
| Sodium Sulfide 400 $\mu$M: nrt1 | −0.119 | 0.297 | −0.402 | 0.688 |
| Sodium Sulfide 400 $\mu$M: dur3 | 0.166 | 0.290 | 0.573 | 0.567 |
| Sodium Sulfide 400 $\mu$M: alr2 | 0.097 | 0.298 | 0.324 | 0.746 |
| Sodium Sulfide 400 $\mu$M: sac3 | 0.390 | 0.539 | 0.725 | 0.469 |
| yhl008c | −3.002 | 0.188 | −15.986 | <0.001 |
| u0394tmn3 | −0.773 | 0.207 | −3.730 | <0.001 |
| put4 | −2.021 | 0.226 | −8.941 | <0.001 |
| pma2 | −0.872 | 0.208 | −4.196 | <0.001 |
| nrt1 | −0.981 | 0.209 | −4.698 | <0.001 |
| dur3 | −0.187 | 0.206 | −0.907 | 0.365 |
| alr2 | −1.145 | 0.211 | −5.431 | <0.001 |
| sac3 | −4.132 | 0.341 | −12.136 | <0.001 |
| 75 $\mu$M Ammonium Sulfate | −0.485 | 0.121 | −3.999 | <0.001 |
| Day 6 | 1.145 | 0.112 | 10.249 | <0.001 |
| 400 $\mu$M Sodium Sulfide | −0.262 | 0.146 | −1.798 | 0.072 |
| Intercept | −0.160 | 0.129 | −1.236 | 0.217 |
| $\phi$ | 5.739 | 0.377 | 15.219 | <0.001 |
| Num.Obs. | 460 | | | |
| R2 | 0.595 | | | |
| R2 Pseudo | 0.595 | | | |
| AIC | −741.6 | | | |
| BIC | −650.7 | | | |
| Log.Lik. | 392.807 | | | |
| RMSE | 0.15 | | | |
| DF Null | 458 | | | |
| DF Resid | 438 | | | |

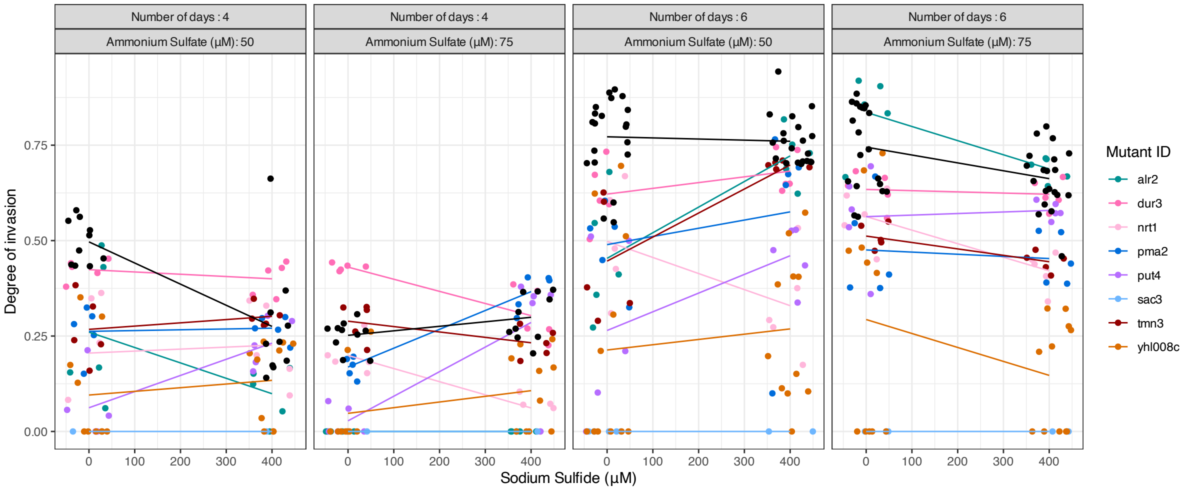


**Figure 4.** Measurement of yeast invasion using image analysis (top row). The x-axis represents the amount of sodium sulfide present, and hence the line represents the difference between no-sulfur and sulfur conditions for each yeast mutant in each experimental condition (color, Parent yeast represented in black). The line is draw between the average values of yeast image for that particular mutant in the experimental condition. In condition on day 6 with higher ammonium sulfate and day 6 the difference between sodium sulfide and no sodium sulfide was as strong as previously observed, making any deviations from this harder to identify.
